# A vector calculus for neural computation in the cerebellum

**DOI:** 10.1101/2024.11.14.623565

**Authors:** Mohammad Amin Fakharian, Alden M. Shoup, Paul Hage, Hisham Y. Elseweifi, Reza Shadmehr

## Abstract

Null space theory predicts that a neuron will often generate spikes not to produce behavior, but to prevent another neuron’s impact on behavior. Here, we present a direct test of this theory in the brain. In the marmoset cerebellum, spike-triggered averaging identified a vector for each Purkinje cell (P-cell) along which its spikes displaced the eyes. Two spikes in two different P-cells produced superposition of their vectors. In the resulting population activity, the spikes were canceled if their contributions were perpendicular to the intended movement. Mossy fibers provided a copy of the motor commands and the sensory goal of the movement. Molecular layer interneurons transformed these inputs so that the P-cell population predicted when the movement had reached the goal and should be stopped.

## Main Text

In order to understand the neuronal computations performed by the brain, current theories often assume the existence of a null space (*1*). Null space is a theoretical concept that was developed to solve a puzzle: why are neurons in the motor regions of the cerebral cortex active not just during production of a movement, but also during its preparation and after its conclusion? The theory assumes that each neuron contributes to a movement with a single vector of weights that represents the effective connection strength of that neuron to the various muscles (*2*). A neuron may be active, but this activity does not necessarily cause a movement because the weighted combination of spikes in the population produces a sum that is zero during certain periods, resulting in cancellation. Thus, the critical element of the theory is the weight vector that is assigned to each neuron.

The problem is that these vectors are currently found through a model that maps the neuronal activities to movement kinematics (*2, 3*). But how can we know what aspect of the movement the neurons are contributing to? That is, how can we test the idea that in the brain, one neuron engages in apparently wasteful production of spikes despite the fact that the downstream effects of those spikes will be nullified by the spikes of another neuron? Indeed, why are the neurons producing these wasteful spikes in the first place?

If each weight vector is a reflection of functional anatomy, then two predictions must hold: 1) when each neuron’s weight vector is estimated not via model fitting, but independently via spike-triggered averaging, then the resulting sum of vectors in the population should cancel before and after the movement, but not during the movement, and 2) simultaneous spikes in two neurons must obey the principle of superposition, as predicted by their weight vectors.

Here, we consider a network of neurons in a region of the cerebellum concerned with eye movements (Fig. S1). This network receives inputs via mossy fibers (MFs) and climbing fibers, and produces outputs via Purkinje cells (P-cells). We first find the weight vector for each P-cell not by fitting its simple spikes (SS) to behavior, but by relying on its climbing fiber input, which produces a stochastic, low frequency (1 Hz) complex spike (CS) that suppresses the SSs completely. We use these *a priori* weights to map the computation in the network from its MF inputs, to the molecular layer interneurons (MLIs), and finally to the P-cell outputs.

### Spike interactions organized neurons in the cerebellar cortex into discrete populations

We trained marmosets to make saccades to visual targets (Fig. 1A) and used Neuropixels and other silicon probes to record from 520 definitive P-cells, 798 putative MLIs (pMLIs), and 1716 putative mossy fibers (pMFs) from lobules VI and VII of the vermis (Figs. S2, S3, & S4). Fig. 1C presents an example session in which we simultaneously isolated over 100 neurons. We identified the P-cells based on their simple and complex spikes and then used the distinct shapes of the CS and SS waveforms in the dendritic tree, soma, and axon of each P-cell (*4, 5*) to identify the molecular, P-cell, and granular layers, respectively.

**Fig. 1.**
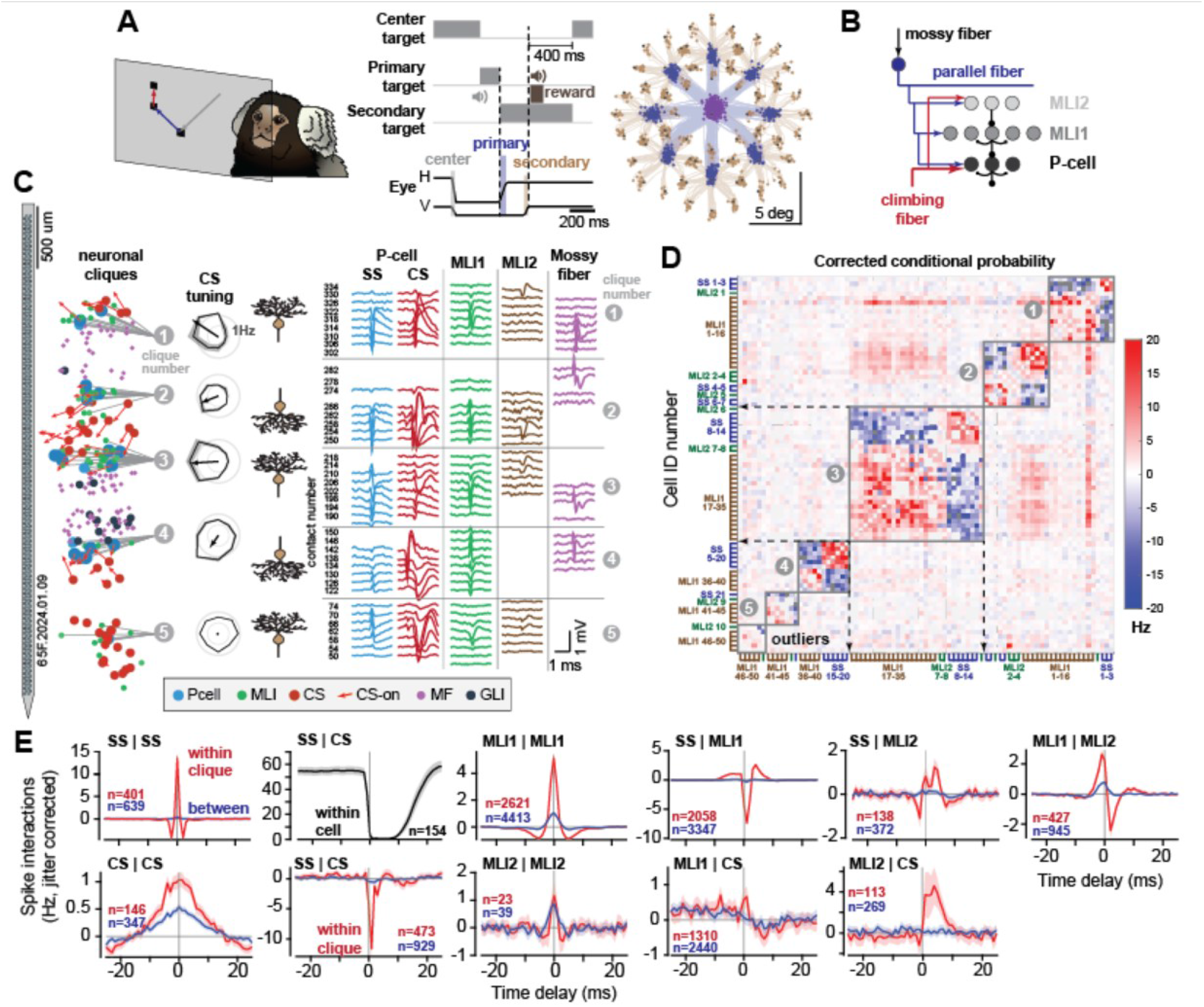
Spike interactions clustered neurons in the cerebellar cortex into discrete populations. **(A)** Random corrective saccade paradigm. **(B)** Architecture of the neural network, with MLI2 inhibiting MLI1, and MLI1 inhibiting P-cells. **(C)** Example Neuropixels session with 5 neuronal cliques. Spike waveforms are displayed for the example neurons. P-cell implies isolation of both SS and CS. CS tuning indicates the average response in the limbing fiber inputs to each clique as a function of direction of visuomotor events. P-cell dendrite and axon orientations are drawn at approximate location of the molecular and granular layers, respectively. **(D)** Adjacency matrix. Rows and columns are neurons recorded simultaneously, with the color intensity of the cell indicating the strength of between-neuron spike interactions (jitter-corrected conditional probability). Cliques and outliers are identified. **(E)** The jitter-corrected probability that a spike occurred in one neuron, given that a spike had occurred in another neuron at a given time delay, for neuron pairs within and between cliques, except for SS|CS, which indicates within neuron suppression of SS following arrival of a CS. Error bars are 2xSEM.

The probe recorded from multiple folia simultaneously, allowing us to ask whether the various neurons clustered into “cliques,” i.e., groups of neurons that exhibited strong spike interactions with each other but not with the other simultaneously recorded neurons (*6*). To quantify these interactions, we computed the jitter corrected (*7*) conditional probability that one cell produced a spike at time *t* + Δ, given that another cell fired at time *t* (Fig. 1E). We then formed an adjacency matrix (Fig. 1D) where the numerical value in each element of the matrix was the strength of the spike interaction between the two cells. We applied graph spectral clustering to the values in the matrix and identified boundaries that divided the neurons into cliques. Clique membership was largely but not exclusively due to the folium boundary.

If two P-cells belonged to the same clique, then their SSs strongly synchronized: an SS in one P-cell accompanied ~14 Hz increase in the SS firing rate of another P-cell at sub-millisecond delay, then a ~5 Hz decrease in firing rate at 3 ms delay (Fig. 1E). This synchrony was absent when the P-cells belonged to different cliques. Similarly, if a P-cell and a pMLI1 were in the same clique, a spike in the pMLI1 produced a reduction of ~7 Hz in the firing rate of the P-cell at 1 ms delay (Fig. 1E). In comparison, there was a reduction of 0.3 Hz when the two cells were from disparate cliques. If the pMLI2 and pMLI1 were in the same cliques, a spike in the pMLI2 produced an inhibition of ~2 Hz in the pMLI1, but little or no inhibition if the two were in different cliques (within vs. between, mean±SEM, SS|SS: 13.6±0.5 vs. 0.4±0.04 Hz, p<1e-100, MLI|MLI: 5.2±0.11 vs. 1.04±0.02, p<1e-100, MLI2|MLI2: 1.17±0.26 vs. 0.86±0.1, p=2e-1, CS|CS: 1.04±0.07 vs. 0.54±0.03, p<1e-12, SS|MLI1 @1ms: −7.38±0.17 vs. −0.26±0.03, p<1e-100, MLI1|MLI2 @2ms: −2.43±0.23 vs. 0.14±0.04, p<1e-50, SS|MLI2 @3ms: 1.24±0.19 vs. 0.12±0.06. p<1e-11). Finally, the climbing fiber input produced a spillover effect upon the pMLI2s (*8, 9*), as well as an ephaptic coupling suppression on the neighboring P-cells (*5*), but only if these neurons belonged to the same clique as the climbing fiber (Fig. 1E) (within vs. between, mean±SEM, MLI1|CS @0-10ms: −0.08±0.04 vs. −0.09±0.02 Hz, p=7.6e-1, MLI2|CS@0-10ms: 2.53±0.43 vs. 0.13±0.05Hz, p<1e-14, SS|CS@1ms: −11.66±0.53 vs. −0.58±0.19Hz, p<1e-100).

We next asked whether clique membership also stratified the neurons based on the information that they received in their inputs via the climbing fibers. To quantify this input, we relied on the fact that following presentation of a visual target at a random direction, the climbing fibers informed the P-cells regarding the direction of that visual event (*10*–*14*) (Fig. 2A). In addition, when a movement was about to be made, the climbing fibers reported the direction of the upcoming movement (*14*). For each P-cell *i*, the climbing fiber encoding of the visuomotor events was along a preferred axis: the CS rates were maximum when the event was in direction *θ*_*i*_ and minimum when the event was in direction *θ*_*i*_ + *π* (Fig. 2A, right plot).

**Fig. 2.**
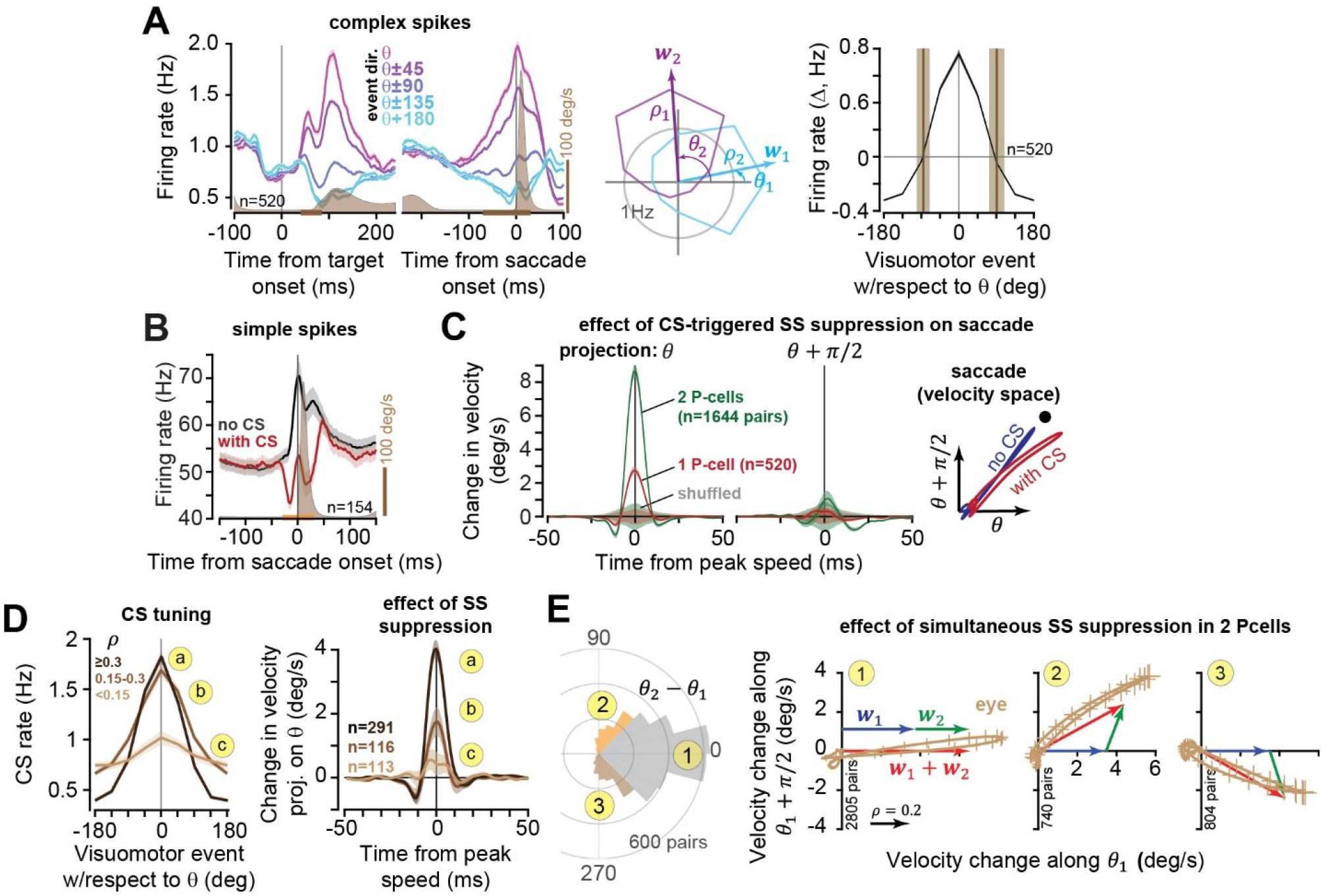
CS-triggered suppression defined a potent vector for each P-cell. **(A)** CS response aligned to the direction of visual and motor events. For each P-cell, the CS response defined a weight vector with amplitude *ρ* and direction *θ*. **(B)** SS rates during saccades that did or did not experience a CS during the period indicated by the orange line. **(C)** Effects of the CS-triggered suppression in one P-cell, or two simultaneously recorded P-cells, on saccade trajectory. Here, pairs of P-cells had |Δ*θ*_*ij*_ | < 22.5 deg. Shaded regions shows 95% confidence intervals for the null hypothesis. **(D)** P-cells were divided into 3 groups (a, b, and c) based on the strength of their climbing fiber input *ρ*. The effect of SS suppression on saccade trajectory was largest in the group that had the largest *ρ*. **(E)** Test of superposition. Pairs of simultaneously recorded P-cells were divided into three groups based on Δ*θ*. We measured effects of simultaneous SS suppression on saccade trajectory. For example, in group 1, the two P-cells had vectors ***w***_1_ and ***w***_2_ that were, on average, parallel. The sum of the two vectors is shown in red, along with the CS-triggered change in saccade trajectory (brown line and error bars). Error bars are SEM.

The strength with which the climbing fibers transmitted visuomotor events differed among the cliques. In some cliques, the climbing fiber inputs exhibited a strong tuning (Fig. 1C clique 1), while for other cliques this tuning was weak (Fig. 1C, clique 5). We represented the information content in each climbing fiber as a vector by fitting a von Mises distribution to the CS rates as a function of direction of the visuomotor event (Fig. 2A, right plot). This produced a vector with direction *θ*_*i*_ and amplitude *ρ*_*i*_ for P-cell *i*. Within a clique, these climbing fiber vectors were more similar to each other than between cliques (Fig. S5, median±MAD: within: 23.25±17.22 deg, between: 57.73±36.05deg, rank sum test p<1e- 17; angle difference from center, mean±std: −1.39±29.99 deg p=0.57). However, while there was strong SS synchrony within a clique, there was little or no synchrony between cliques, even when they received similar climbing fiber information (Fig. S6). Therefore, cliques defined a smaller unit of computation in the cerebellum than climbing fiber defined microzones (*15*).

### The climbing fiber input identified a potent vector for each P-cell

The climbing fiber input completely suppressed the P-cell’s ability to produce simple spikes (Fig. 1E, Fig. S3A). When this suppression took place during a movement, it slightly diverted that movement, acting as a stochastic perturbation (*16*–*18*). We performed spike-triggered averaging, comparing saccades that had the same starting position and were made toward the same target, but did or did not experience a CS-triggered SS suppression (Fig. 2B). We evaluated the difference in the eye trajectories in velocity pace. This difference produced a vector that throughout the duration of the saccade remained parallel to *θ*_*i*_ (Fig. 2C, Fig. S7). When two P-cells with similar *θ* were suppressed during the same saccade, the amplitude of the disturbance more than doubled. Thus, when P-cell *i* was suppressed via its climbing fiber input, the result was a displacement of the eyes along a vector that was on average parallel to direction *θ*_*i*_.

For each P-cell, represent *θ*_*i*_ as 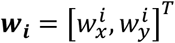. To assign a magnitude to this vector, we considered the possibility that suppression of the P-cells that belonged to a clique with a strong CS tuning (i.e., large *ρ*) produced a greater pull on the eyes, as compared to other cliques that had a weak CS tuning (Fig. 2D). The greater the amplitude *ρ*, the greater the displacement caused by the SS suppression (Fig. 2D, right subplot, projection on *θ*, ANOVA F(2)= 27.3, p = 5.3e-12). We assigned the following vector to each P-cell and labeled it as its potent vector:

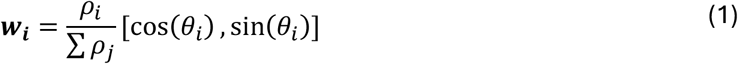

Thus, P-cell *i* was assigned a potent vector with direction *θ*_*i*_ and amplitude 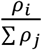.

### P-cells obeyed superposition in the space of their potent vectors

We next asked whether the downstream effect of suppressing two P-cells simultaneously was a linear sum of their potent vectors. This was a test of superposition: when P-cells *i* and *j* were suppressed, was the resulting displacement described by the vector ***w***_***i***_ + ***w***_***j***_? For all pairs of simultaneously recorded P-cells, we selected pairs with 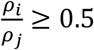, then measured the difference in the angles of the two potent vectors ***w***_***i***_ and ***w***_***j***_, i.e., Δ*θ*_*ij*_ = *θ*_*i*_ − *θ*_*j*_, then used the resulting distribution of Δ*θ*_*ij*_ to divide the pairs into three groups (Fig. 2E). For example, in group 1, Δ*θ*_*ij*_ ≈ 0, i.e., ***w***_***i***_ and ***w***_***j***_ were approximately parallel, whereas for group 2, Δ*θ*_*ij*_ ≈ 70 deg, i.e., ***w***_***i***_ and ***w***_***j***_ were on average 70 degrees apart. Indeed, simultaneous suppression of two P-cells pulled the eyes along a vector with magnitude and direction that was predicted by ***w***_***i***_ + ***w***_***j***_ (red vector, Fig. 2E), demonstrating superposition. Because a single climbing fiber innervates ~8 P-cells (*19*), and there is CS synchrony within a clique (Fig. 1E), these effects are likely due to simultaneous suppression of many P-cells.

### The potent vector defined the axis of symmetry for the simple spikes

Some P-cells increased their SS rates around saccade onset, while others decreased their rates (Fig. 3A, top row). Given that the downstream effect of SS suppression in both groups was to pull the eyes in direction *θ* (Fig. S7B) (*17*), it was puzzling that the modulations were present for all saccade directions and continued long after the movement had ended.

**Fig. 3.**
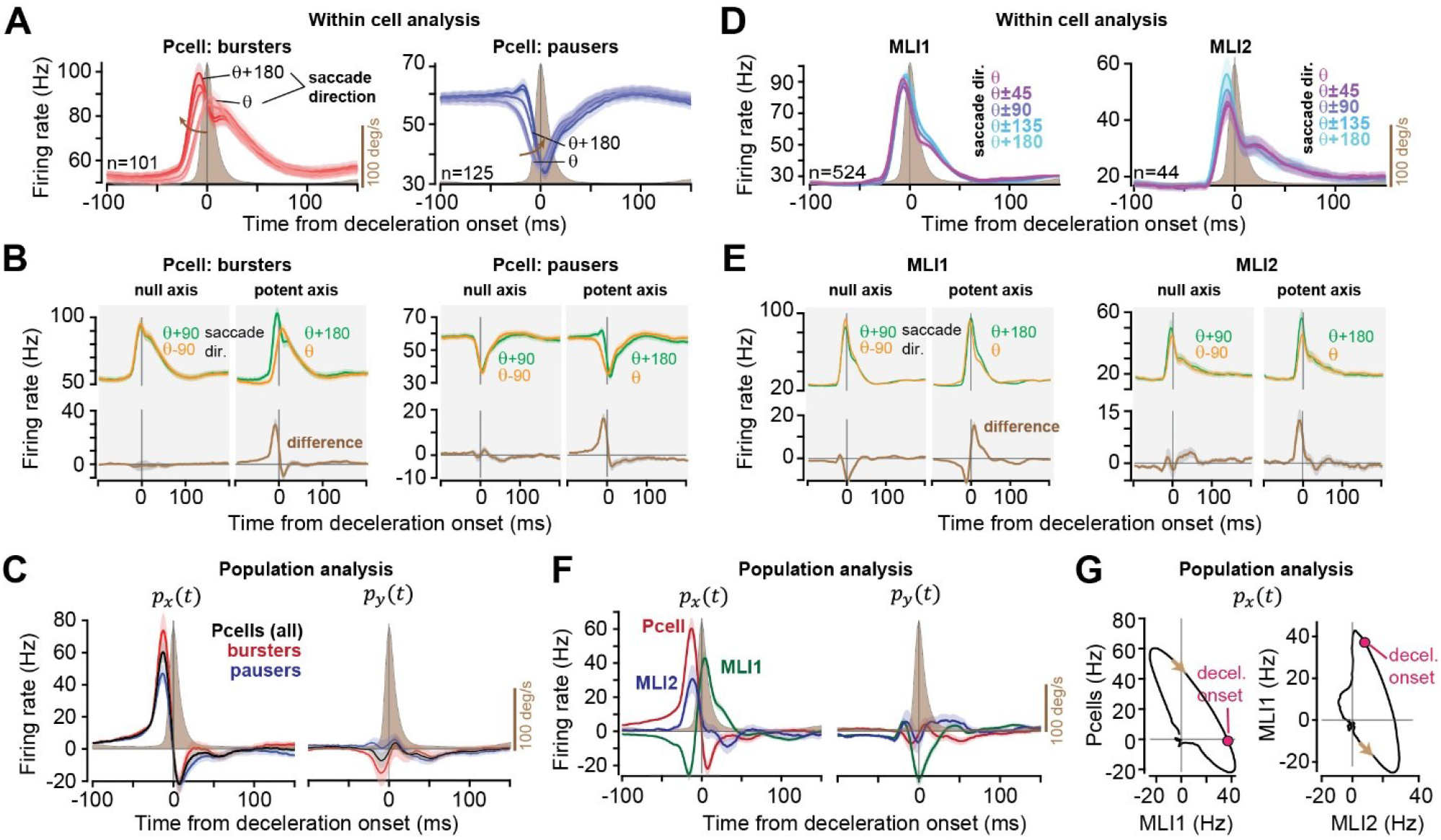
Neurons within a clique had symmetric spike rates with respect to the potent axis, canceling the spikes that had downstream effects perpendicular to the direction of the movement. **(A)** Average P-cell response (red: bursters, blue: pausers) to saccade direction with respect to potent vector of each neuron, *ψ* − *θ*. **(B)** SS rates for saccades in direction 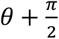 are equal to the rates for saccades in direction 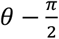. In contrast, the rates for saccades parallel to the potent vector are not equal. **(C)** Population response of all P-cells, i.e., the sum of rates, weighted by each neuron’s 2D potent vector. **(D)** Same as A for the pMLIs. **(E)** Same as B for the pMLIs. **(F)** Population response of the pMLIs. **(G)** Phase plots of P-cell potent response as a function of pMLI1 (left), and pMLI1 as a function of pMLI2 (right). The negative slope indicates that in the population response, there is a negative correlation between the firing rates of the pMLI1s and the P-cells, and pMLI2s and pMLI1s. Error bars are SEM.

However, if we represented the SS rates *r*_*i*_(*t*) as a function of saccade direction *ψ* with respect to the direction of the potent vector *θ*, then a remarkable pattern emerged: when the saccade was in direction 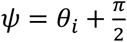, the SS rates *r* (*ψ, t*) in the population were nearly identical to when the saccade was in direction 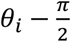. As a result, a within P-cell subtraction of the rates in these two directions produced, on average, complete cancellation (Fig. 3B, bottom row). This pattern held true for pairs of saccades that were at an equal angle with respect to the potent axis *θ* (Fig. S8). Thus, we made the following inference regarding the within-cell symmetry in the simple spikes:

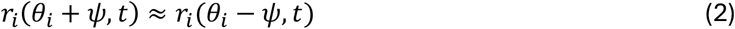

Eq. (2) held true in both the burster and the pauser P-cells. Thus, just as the CS rates were, on average, symmetric with respect to *θ* (Fig. 2D) (a finding that is consistent in both marmosets and macaques (*11*)), so were the SS rates.

In contrast, when one saccade was in direction *ψ* = *θ*_*i*_ and another along *ψ* = *θ*_*i*_ + *π*, now the within-neuron difference in the SS rates produced a burst-pause pattern that crossed zero at the deceleration onset of the saccade (Fig. 3B, second column). This pattern held true in the bursters as well as the pausers, despite the fact that the pausers never produced an increase in their activity. Moreover, while the SS rates for saccades in all directions in both the bursters and pausers had remained modulated long after the movement had ended, in both groups the extra spikes were canceled for saccades parallel to the potent vector, producing a burst-pause pattern that returned to zero as the saccade ended (Fig. 3B, S17, S18).

### The potent vector defined a population code for the simple spikes

According to null space theory, a neuron’s contribution to a movement is described by a constant weight, which is usually found via the parameters of an equation that maps firing rates to behavior (*2*). Here, rather than fitting such a model, we had the weights for mapping the simple spikes from the complex spikes: when a movement was made in direction *ψ*, P-cell *i* produced simple spikes with the rate *r*_*i*_(*ψ, t*), and its contribution to behavior was defined by its potent vector ***w***_***i***_:

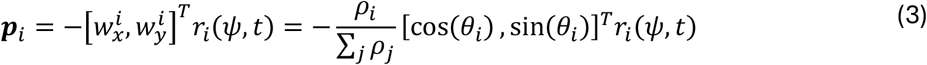

The negative sign is because suppression of SS produced forces in direction *θ*_*i*_. Because *θ*_*i*_ depended on the neuron’s anatomical location in the cerebellum (*11, 14*), we removed our sampling bias (Fig. S6B) by assuming isotropy for *θ*, duplicating each P-cell uniformly as 8 P-cells:

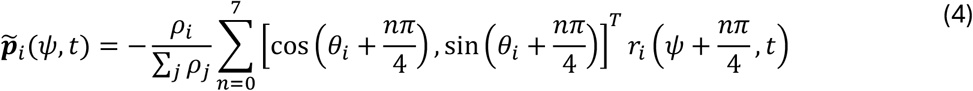

The output of the population was simply the weighted sum of activity in all neurons:

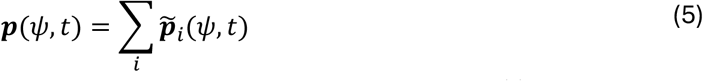

Without loss of generality, consider a movement in direction *ψ* = 0. The vector ***p***(*t*) has two components: *p*_*x*_(*t*), i.e., the SS rates that move the eyes along the direction of the movement, and *p*_*y*_(*t*), i.e., the SS rates that move the eyes perpendicular to it. We found that *p*_*x*_(*t*) exhibited a positive-negative pattern that crossed zero at the onset of the saccade’s deceleration (Fig. 3C, all P-cells, left panel). In contrast, *p*_*y*_(*t*) remained near zero throughout the movement. Thus, despite the fact that in the individual neurons the SS rates were modulated for saccades in all directions, in the population the downstream effects of the SSs were canceled so that they had no effects perpendicular to the direction of the movement (Fig. 3C, right panel). Even though the SSs for individual P-cells were modulated long after the saccade ended, the downstream effects were canceled beyond the end of the movement (Fig. 3C, left panel, Fig. S17). Finally, despite the fact that some P-cells exhibited a burst, while others exhibited a pause, in both groups *p*_*x*_(*t*) exhibited a positive-negative pattern that crossed zero at deceleration onset, and in both groups *p*_*y*_(*t*) remained near zero throughout the movement (Fig. 3C).

### A P-cell’s contribution to behavior was defined primarily by the magnitude of its potent vector, not the magnitude of its SS modulation

It is natural to assume that when a neuron’s firing rates are strongly modulated during a behavior, it is contributing significantly to the control of that behavior. However, our theoretical framework makes a counter-intuitive prediction: because the downstream effects of the spikes in one neuron may be canceled by another neuron, what matters is not how much the SS rates were modulated, but how large was the potent vector. For example, consider two groups of P-cells that had roughly equal SS rate modulation during saccades, but different *ρ*: one group had a large *ρ* (i.e., large amplitude potent vector), the other a small *ρ* (Fig. S9A). For each P-cell, *ρ* indicates the sensitivity of the CS rates for sensorimotor events in directions *θ* vs. *θ* + *π*. Because the complex spikes are long-term teachers that guide sculpting of the simple spikes (*12, 20*), a large *ρ* implies that the teacher dissociates between the directions of these sensorimotor events, which implies that the SS rates should be different. Therefore, the theory predicts that in the population, for P-cells that have a small *ρ*, the SS rates *r*(*θ, t*) will be similar to *r*(*θ* + *π, t*), resulting in greater cancellation than for P-cells that have a large *ρ* (Fig. S18).

To test this prediction, in our two groups of larger and smaller *ρ* P-cells, we set *ρ* = 1 for all cells and then computed the population response via Eq. (5). Whereas the two groups had roughly equal SS rate modulations as a function of movement direction, the population response of the large potent vector group was roughly twice as large as the small potent vector group (Fig. S9B). Thus, among the P-cells with small potent vectors, despite their large modulation of simple spikes during saccades, most of these spikes were canceled.

### The interneurons inherited their potent vector from the P-cells

In both types of MLIs, the rates in all saccade directions exhibited a burst (Fig. 3D), which was difficult to understand in the context of anatomy: after all, pMLI2s inhibited pMLI1s, and pMLI1s inhibited the P-cells. Why were the interneurons bursting in all directions?

Each clique had P-cells with a potent vector ***w***. We used this vector to analyze the firing patterns of the MLIs. The potent axis of the P-cells was inherited by the MLIs such that the axis of symmetry roughly held true for the interneurons. When one saccade was in direction *ψ* = *θ* and another in direction *ψ* = *θ* + *π*, the within-cell difference in the rates was a pause-burst pattern in the pMLI1s, and a burst pattern in pMLI2s (Fig. 3E). When the two saccades were in directions *ψ* = *θ* + *π*/2 and *ψ* = *θ* − *π*/2, the difference in the two rates was nearly zero for pMLI2s and missing the burst in pMLI1s.

To estimate the population response of the pMLIs, we applied the same analysis as for the P-cells: we used Eqs. (4) and (5) and assigned to each pMLI the potent vector of its clique. Now we found that in the population of pMLI1s, in the direction of the target the response was a pause-burst pattern (Fig. 3F, left panel). In the pMLI2s, the population response was a burst that returned to zero at deceleration onset. Plotting these population rates of the various neurons against one another (Fig. 3G) unmasked a negative correlation, consistent with the inhibitory influence that the pMLI2s have on pMLI1s, and the inhibitory influence that the pMLI1s have on the P-cells.

The population response in pMLI2s perpendicular to the target remained nearly zero during the saccade, exhibiting cancellation (Fig. 3F, right panel). However, this was not the case for pMLI1s, an inconsistency that we do not currently understand, but may be related to the fact that the pMLI1s appear to be composed of two distinct subtypes (*21*).

### The potent vectors revealed the computations performed in this region of the cerebellum

The burst-pause pattern in the P-cells implied a computation associated with predicting when to stop the saccade. For this to be possible, this region of the cerebellum should receive two kinds of information: a copy of the motor commands in muscle coordinates, and a copy of the goal location in visual coordinates. To examine this hypothesis, we recorded from n=1716 pMFs (Fig. S10) and found one group of inputs that provided information regarding the goal of the movement in visual coordinates, termed goal pMF (Fig. 4A, Fig. S11), and another group that encoded the motor commands, termed state pMF (Fig. 4D, S12, S13).

**Fig. 4.**
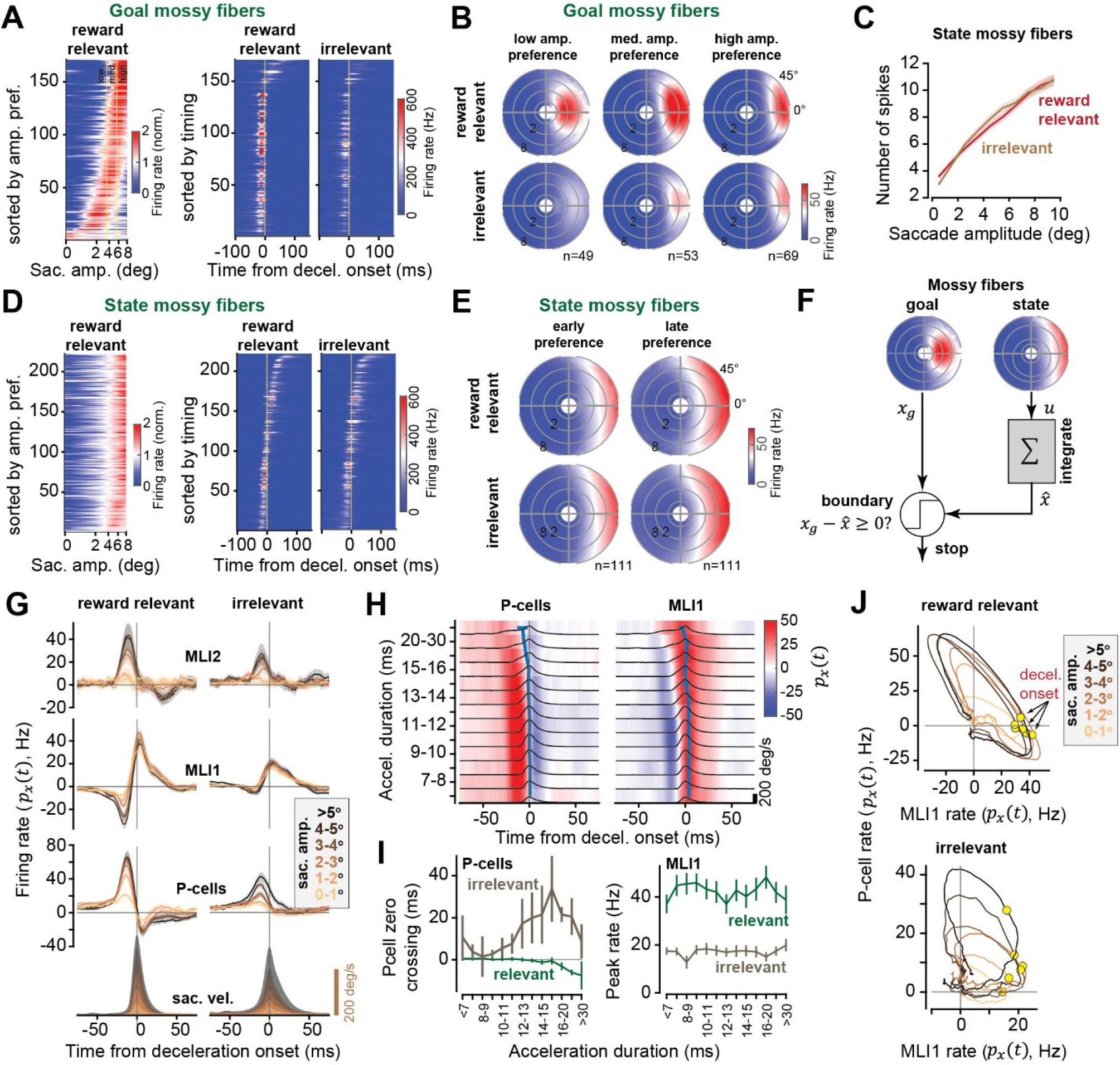
P-cells signaled when the saccade should end, but only if the mossy fibers provide goal location information. Two groups of pMFs carried information regarding the sensory goal location and the motor commands (goal and state MFs). **(A)** Left, Normalized responses of goal pMFs to different saccade amplitudes in the preferred direction. Right, average responses of the same pMFs aligned to deceleration onset for reward relevant and irrelevant saccades (for saccades with similar peak velocities, 150-250°/s). **(B)** Polar plots presenting the movement fields of the goal pMFs as a function of amplitude preference. **(C)** Number of spikes during the saccade as a function of saccade amplitude for state pMFs. **(D-E)** Same as A-B for state pMFs. **(F)** A model: an integration of motor commands estimates the displacement and a comparison between the displacement and the goal can signal when to stop the saccade. **(G)** Clique pMLIs and P-cell responses along the potent axis for relevant and irrelevant saccades binned by peak velocity. **(H)** P-cell and pMLI1 potent responses for relevant saccades binned by acceleration duration. **(I)** P-cell zero-crossing and pMLI1 peak rate for different acceleration durations. **(J)** P-cell potent response as a function of pMLI1 potent response for reward relevant (*top*) and irrelevant (*bottom*) saccades. In spite of negative correlation in both cases, deceleration onset is indicated at a specific phase only for reward relevant saccades. Error bars are SEM.

The goal pMFs generally exhibited buildup activity before saccade onset with a relatively narrow spatial response that encoded the location of the visual target with respect to the fovea (Fig. 4B, S13E-F). However, the goal information was present only if a reward relevant visual target was the destination of the saccade. When a similar saccade was made spontaneously without this target, the goal pMFs exhibited a muted response (Fig. 4A-B, S11B-E). In contrast, the state pMFs continued to provide a faithful copy of the motor commands regardless of whether the movement was reward relevant or not (Fig. 4D-E, S12E-F). Indeed, a simple integration of the number of spikes in the state pMFs was an excellent predictor of the saccade amplitude in all conditions (Fig. 4C).

These results suggested a working model (Fig. 4F). When the saccade was aimed at a reward relevant target, the cerebellar cortex was made aware of the target location *x*_*g*_ via the goal pMFs. We speculate that the network, possibly relying on unipolar brush cells in the granular layer (*22*), integrated the spikes in the state pMFs to a bound set by the goal pMFs. Once the bound was reached, the P-cells signaled to the nucleus via disinhibition to produce forces that would aid in stopping the movement. The model predicted that when the goal information was withheld from the cerebellum (i.e., reward irrelevant saccades), the integration would still occur, but the P-cells would not be able to provide a stopping signal.

To test this prediction, we binned the movements based on saccade peak velocity and acceleration durations and found that regardless of saccade velocity, amplitude, or acceleration duration, in reward relevant saccades the SS rates *p*_*x*_(*t*) transitioned from burst to pause at deceleration onset (Fig. 4G, first column, Fig. 4H & 4I, Fig. S14). The one exception was for very long saccade acceleration periods. In these saccades, the velocity during the acceleration phase was double-peaked, and both the goal and state pMFs showed a double peak in their rate profiles, indicating rare saccades in which mid-flight, the goal was changed (Fig. S15). For these “double saccades”, the SS rates crossed zero before deceleration onset, but for all other reward relevant saccades the zero-crossing remained locked to deceleration onset. Moreover, regardless of saccade velocity, amplitude, or acceleration duration, if the target was reward relevant then the firing rates of pMLI1s reached a constant peak value (Fig. 4H-J).

These features were missing for the irrelevant saccades. For example, in the irrelevant saccades the SS rates exhibited an increase as a function of velocity but did not cross zero (Fig. 4G, S14). Also, the firing rates of pMLI1s reached half the peak for reward irrelevant saccades (Fig. 4G, S14).

Finally, we considered the spikes that moved the eyes perpendicular to the direction of the target and found that if the movement was in the context of reward, spike cancellation became more precise as saccade amplitude and velocity increased (Fig. S16). In contrast, when no reward was at stake, the opposite happened; spike cancellation degraded as the movement’s amplitude and velocity increased.

## Discussion

We uncovered the logic of computation in a region of the cerebellum that is concerned with control of eye movements, finding evidence that neurons often generated spikes not to produce behavior, but to prevent the effects that other neurons would have on behavior.

The P-cells, the interneurons, and the climbing fiber inputs exhibited spike interactions that grouped them into discrete networks, called cliques. Using the climbing fiber input, we found that each P-cell pulled the eyes along a specific 2D vector, called the potent vector. However, the P-cells and the interneurons were modulated not just for saccades along their potent vector, but for all saccades. Indeed, their modulation continued long after the movement had ended. Critically, when two P-cells with different potent vectors were simultaneously suppressed, the downstream effect on behavior was a summation of the individual vectors. This raised the possibility that some neurons were modulated not to contribute to a movement, but to eliminate another neuron’s unwanted contributions to that movement.

A critical clue emerged in the symmetry that was present in the P-cell SS rates: the modulations were nearly equal when two saccades had the same absolute value of angular displacement with respect to the cell’s potent vector. The purpose of this symmetry was revealed in the population. As each P-cell generated its simple spikes during a saccade and consequently pulled the eyes along its potent vector, the superposition of the potent vectors in the population removed the component that would have pulled the eyes perpendicular to the direction of movement. What remained was a vector in the direction of the movement, exhibiting a fast-changing burst-pause pattern that consistently crossed zero at the onset of deceleration. The same potent vector explained how the interneurons in the clique produced spikes that could guide the output of the P-cells. Thus, the P-cells as a population produced an output that assisted with both initial acceleration and final deceleration.

A neural circuit that produces such an output must have access to two pieces of information: the goal location in sensory coordinates *x*_*g*_, and the real-time commands in motor coordinates *u*(*t*) (Fig. 4F). The mossy fiber inputs separately provided each piece of information, but only if the planned movement was reward relevant. For the irrelevant saccades, the mossy fibers continued to provide an accurate copy of the motor commands, but the goal information was muted, producing a population output in the P-cells that was a burst without a pause. As a result, when reward was not at stake, the cerebellum received degraded information about the goal of the movement, the P-cells were unable to predict when to stop that movement, and the spikes that would divert the movement away from the intended target were poorly canceled.

Thus, the population output relied on competitive spike cancellation among the neurons, an idea that was first introduced for P-cells in (*11*). That work demonstrated cancellation as a function of time when saccades were in the preferred direction of the climbing fiber input. Here, we extended that theory by establishing the concept of a potent vector for the P-cells as well as the interneurons, discovering complete cancellation of spikes among the P-cells when movements were perpendicular to their potent vector.

But why should a neuron produce spikes if its downstream consequences are nulled by another neuron? The answer appears to be in the difference between what the population produced, and what the individual neurons produced (Fig. S17): because the population output was a fast-changing function of time, the much slower individual P-cells appeared to be placed in subtractive competition with each other, eliminating each other’s spikes to produce the fast dynamics of the output. The same pattern was present in the pMLIs: their population response exhibited a fast changing pause-burst output, despite the fact that all the pMLIs were bursters.

However, this explanation alone is not sufficient to account for all our data. All the SS modulations produced by one group of P-cells for saccades in directions perpendicular to their potent vector were canceled by another group of P-cells with the opposite potent vector (Fig. S20). The rationale for a neuron to produce spikes despite complete cancellation by another neuron remains unknown, but perhaps is indicative of a reserve for learning, for example, in case there is unilateral weakness in the extraocular muscles (*23*). A more likely explanation is that gaze control normally includes eye, head, and body movements, and the complete cancellation that we observed here would not occur if other body parts were allowed to move and thus contribute to directing the gaze.

Cerebellar disease often produces loss of P-cells, for which there are few therapeutic options. Here, we discovered a vector calculus that maps the mossy fiber inputs in this region of the cerebellum to the P-cell outputs via the MLIs. Our theory can contribute to building of circuits that emulate the neural computation of a healthy cerebellum and functionally replace the damaged tissue (*24*).

## Acknowledgements

We are thankful to S.Z. Muller for discussions, T. Harris and A. Graves for Neuropixels Ultra probes and the hardware to use them, X. Wang for the marmoset colony, and J. Izzi, the lead veterinarian.

## Funding

National Institutes of Health grants NS128416 (R.S.) and EB028156 (R.S.).

## Author contributions

Methodology: MAF, AMS, PH, HYE, RS

Investigation: MAF, AMS, PH

Visualization: MAF, HYE

Writing: RS and MAF

## Competing interests

The authors declare that they have no competing interests.

## Data availability

All data and code are available at: https://doi.org/10.5061/dryad.qnk98sft9

## Supplementary Materials

Materials and Methods

Figs. S1 to S21

References (25-67)

## Notes

https://doi.org/10.5061/dryad.qnk98sft9

### Competing Interest Statement

The authors have declared no competing interest.

### Summary of Updates

1.We added a new figure to provide an overview of the problem that the manuscript solved (Fig. S1). This figure summarizes the steps that we took to understand the computations that are performed by the neurons in a region of the cerebellum. 2.We added new results showing that cliques, as defined via their spike interactions, appear to be a finer unit of computation in the cerebellum than climbing fiber defined microzones (Fig. S6)

