## Supplementary materials for "A vector calculus for neural computation in the cerebellum"

The PDF file includes:

Materials and Methods
Figs. S1 to S21
References

Other Material for this manuscript includes the following:

MDAR Reproducibility Checklist

**Materials and Methods**

Data were collected from two marmosets, *Callithrix Jacchus*, 350-370 g, subjects 132F (Charlie, 7 years old), and 65F (Barney, 7 years old), during a 2-year period. The marmosets were born and raised in a colony that Prof. Xiaoqin Wang has maintained at the Johns Hopkins School of Medicine since 1996. Our procedures were approved by the Johns Hopkins University Animal Care and Use Committee in compliance with the guidelines of the United States National Institutes of Health.

*Data acquisition*

Following recovery from head-post implantation surgery, the animals were trained to make saccades to visual targets and rewarded with a mixture of applesauce and lab diet (*25*). Visual targets were presented on an LCD screen with 500Hz refresh rate and low latency (Dell AW2524H). Binocular eye movements were tracked at 2000 Hz using VPIXX eye tracking system. Tongue movements were tracked with a 522 frame/sec Sony IMX287 FLIR camera, with frames captured at 100 Hz.

We performed MRI and CT imaging on each animal and used this data to design an alignment system that defined trajectories from the burr hole to various locations in the cerebellar vermis (*25*), including points in lobule VI and VII. We used 3D Slicer software to first align the T2 MRI to the marmoset atlas (*26*). We then aligned the CT scan image to the transferred T2 MRI (Fig. S1). We used a piezoelectric, high precision microdrive (0.5 micron resolution) with an integrated absolute encoder (M3-LA-3.4-15 Linear smart stage, New Scale Technologies) to advance the electrode. In order to reach the cerebellar cortex with Neuropixels NHP 1.0 and Neuropixels Ultra probes, we planned a posterior burr hole and avoided the confluence of sinuses using MRI T2 images. This approach enabled us to record from multiple folia in lobules VI and VII simultaneously.

We recorded from the cerebellum using Neuropixels 1.0 NHP probes, Neuropixels Ultra probes, as well as 64-channel checkerboard or linear high-density silicon probes (M1 and M2 probes, Cambridge Neurotech). For Neuropixels we used the National Instrument acquisition system (NI PXIe-1071). Data were sampled at 30 kHz and aligned to the eye tracking system time as the reference time using random TTL signals. For the M1 and M2 probes we connected each electrode to a 64-channel head stage amplifier and digitizer (RHD2132 and RHD2164, Intan Technologies, USA), then connected the head stage to a communication system (RHD2000 Evaluation Board, Intan Technologies, USA). Because the conductive coating on the Cambridge probes degraded after each insertion into the brain, we re-coated the probes and restored their low impedance after every 3-4 recording sessions (*27*). For accurate reaction time and visual target time measurements, we measured the monitor latency in real-time with a photodiode and aligned it to the reference time.

*Behavioral protocol*

Each trial began with fixation of a center target after which a primary target appeared at one of 8 randomly selected directions at a distance of 5-6.5 deg. As the subject made a saccade to this primary target, that target was erased, and a secondary target was presented at a distance of 2.5-3.5 deg, also at one of 8 randomly selected directions. The subject was rewarded if following the primary saccade, it made a corrective saccade to the secondary target, landed within a 1.5-2.0 deg radius of the target center, and maintained fixation for at least 200 ms (Fig. 1A). The reward was food that was provided in two small tubes (4.4 mm diameter), one to the left and the other to the right of the animal. A successful trial produced a food increment in one of the tubes and would continue to do so for 50 consecutive trials, then switch to the other tube. Because the food increment was small, the subjects naturally chose to work for a few consecutive trials, tracking the visual targets and allowing the food to accumulate, then stopped tracking and harvested the food via a licking bout (*18*, *28*).

We analyzed eye movements using a deep neural network that detected all saccades and microsaccades (*29*). The pre-trained networks for human and macaque monkeys did not perform well for marmosets. Hence, we designed a custom Matlab GUI-based program to curate the saccades (https://github.com/ShadmehrLCMC/SACCURATE). We then retrained the network through transfer learning for each individual animal using seven 30-45 minutes of recording. To prevent any potential false positive saccades, we pruned the saccades by fitting a bivariable Gaussian distribution to two different feature spaces defined as biologically relevant metrics (*30*) and removed saccades that were outliers (less than 1% chance of belonging to the distribution). The two feature spaces were the log-log main-sequence plots (maximum velocity vs. amplitude, Fig. S4) and the log-log acceleration time to deceleration time ratio vs. amplitude of saccade. We then analyzed and found valid fixations among candidate fixations after or before each detected saccade. We measured and used data-driven thresholds on steady eye position criteria including maximum displacement, dispersion on x and y axes, fixation duration as well as maximum velocity. We discarded fixation candidates with lost eye signals due to instability of eye tracking or blinking.

We detected the onset and offset time of each saccade using a trained neural network, as described above. We then low pass filtered the eye position traces with a 3rd order Butterworth filter with 100 Hz cut-off frequency. We then calculated the saccade velocity by differentiating the eye position trace and found peak velocity times and values. Using the onset, offset and maximum velocity times we formed the acceleration and deceleration duration of each saccade.

Our data included rare saccades with extremely long acceleration durations that had a double-step velocity profile (Fig. S15). We labeled these as double saccades and analyzed them separately, finding that the goal information provided in the mossy fiber inputs appeared to change mid-saccade.

*Neurophysiological analysis*

We used OpenEphys (*31*) for electrophysiology data acquisition, and then used Kilosort 2.0 and Phy 2.0 (*32*) to manually identify and curate the spikes. Each recording was curated twice by two experienced neurophysiologists. We controlled for contamination in the cross-correlograms in P-cell simple and complex spikes and removed the spikelets and double detection of CSs as SSs by aligning the two waveforms to each other and removing the copies in SS units (*33*). We further tracked the sorted units across three to seven 30-45 minute recordings using a semi-supervised custom written MATLAB GUI software. To do so, we used electrophysiology properties of the cells including waveform and location on the probe, auto-correlogram and raster plots aligned to saccade onsets. We discarded unstable cells with varying baseline firing rates across time (commonly due to the physical pressure of the probe). We performed manual cell type identification through identification of the layers. First, we identified the P-cells via suppression of SS following a CS. As P-cells are large cells, their spike waveforms are present on multiple channels of the silicon probes (Fig. 1C). Next, using the CS waveforms on the channels we identified the orientation of the P-cell's axon and dendrites, thus identifying the molecular, Purkinje, and granular layers (*4*, *5*, *33*).

For example, we identified the molecular layer via the downward, broad CS waveforms in the dendrite tree of the P-cells (channels 58-70, CS traces, Fig. 1C, S3B). In the granular layer we looked for spike waveforms that exhibited an "m" shape with a slow negative after-wave and labeled those cells as pMF glomeruli (*34*–*36*) (Fig. 1C, S10A-E). In the molecular layer, we labeled neurons that inhibited the P-cells at 1ms latency or sooner as putative MLI1s (pMLI1), and neurons that inhibited pMLI1s, excited a P-cell, and experienced excitation from CS spillover as pMLI2s (*8*, *9*, *37*) (Fig. 1D-E).

Quantifying the quality of isolation of each neuron. A summary of the measures used to verify the quality of the neurophysiological data is provided in Fig. S3A. To measure the isolation quality of each neuron, we computed the conditional probability $\Pr(s(t+\Delta)=1|s(t)=1)$, that is, the probability of a spike at time delay Δ, given that the neuron produced a spike at time zero. We then multiplied this probability by 1000 (bin size 1 ms) and plotted the results as firing rate. For well-isolated cells, we expected a clean refractory period. To measure the noise rate in each neuron's spiking, we quantified this conditional probability at $\Delta = 1\text{ms}$, except for complex spikes, for which we used a 5ms period. The data for all cell types are shown in Fig. S3A, second row. For example, to quantify the quality of the P-cells, we measured the noise rate and found that for the simple spikes, this rate was 0.58±0.41 Hz (median ± median absolute deviation, MAD). For the complex spikes, the noise rate was 0±0 Hz.

Identification of the MLIs. Following the identification of the molecular, P-cell, and granule layers, we identified the putative MLI1 and MLI2s (labeled as pMLI1 and pMLI2) based on their spike interactions with each other and the P-cells (*37*). These interactions were extracted and measured through cross-correlograms that were then corrected via spike jittering: for each cell pair, the interaction was measured via a cross-correlation, then corrected for chance interaction due to firing rate fluctuations using interval jittering (*7*), as shown in Fig. 1E. pMLI1s were identified using their milliseconds inhibition of P-cells as well as their synchrony with each other, while pMLI2s were identified through their millisecond suppression of pMLI1s and lack of interaction or later disinhibition of P-cells (*37*). We relied on the interactions of MLIs with P-cells to identify them, then confirmed that their waveform, baseline rates, and auto-correlograms were similar to previous findings in definitively identified neurons in mice (*37*) (Fig. S3).

Identification of the mossy fibers: We identified the mossy fibers based on their unique "m" shaped spike waveform (Fig. 1C) in the granule layer. We did this by relying on our Neuropixels Ultra recordings to acquire a very high resolution "picture" of the mossy fiber spike waveforms (Fig. S10). As previously reported, mossy fibers exhibit a triphasic waveform with a negative after wave or a fast narrow spike (*36*, *38*–*42*). We used an approach similar to (*43*) to extract spatio-temporal waveforms of each cell by concatenating spike waveforms on different channels to form a 2D image, then centered the absolute peak to the center of image in time and space (Fig. S10C). In order to get a consistent result for the checkerboard and linear probes we picked two columns out of four columns of the checkerboard probe with highest waveform values for each cell and linearized it. We then centered the absolute peak of the waveform in time and space (Fig. S10C). Finally, we used UMAP to cluster the mossy fibers from other granular layer interneurons (Fig. S10D). UMAP on a linearized feature space of these spatio-temporal waveforms revealed two groups of mossy fibers and a potential Golgi cell group. As the mossy fibers exhibited different spike waveforms based on the location of recording (Fig. S10A, Neuropixel Ultra recording), we found one UMAP group with downward fast spiking waveform and another UMAP group with upward triphasic waveform with a negative after wave. The GLI group, on the other hand, had a larger signature in space and showed wider spike waveforms (Fig. S10E).

Identifying mossy fibers that encoded goal and state mossy fibers. The oculomotor region of the vermis receives mossy fibers mainly from the pontine nuclei, the nucleus reticularis tegmenti pontis (NRTP), and the paramedian pontine reticular formation (PPRF) (*44*) (Fig. S10F). NRTP neurons receive information regarding the goal of the movement from the superior colliculus (*45*, *46*). The PPRF region houses the horizontal burst generator neurons (*47*), which encode saccade velocity along a preferred direction (horizontal or vertical, associated with the motoneurons that innervate the eye muscles). This suggested that the mossy fibers in this region of the cerebellum should separately encode the goal of the movement, and a copy of the motor commands (Fig. S11-S13).

We measured the firing rates of mossy fibers during fixation, saccades, and after presentation of the visual cue. One group of mossy fibers, termed position MFs, had responses similar to neural integrator neurons in the brainstem, exhibiting tonic rates that encoded position of the eyes (Fig. S13B-C). To identify these neurons, we computed the firing rates during all fixations (>200ms fixation) in a 50ms window centered during the fixation period. We then fitted the projected position of the eye on the preferred direction of the mossy fiber and used R-squared larger than 0.25 to identify fixation encoding mossy fibers. We removed these mossy fibers from the population. Next, we removed mossy fibers with less than 500 targeted saccades toward their preferred direction. We separated the remaining mossy fibers based on their baseline firing rates (higher than 20Hz) because neurons in NRTP and PPRF regions usually have low baseline firing rates. Mossy fibers with higher than 20Hz baseline rates showed heterogeneous eye-related activity during saccades (Fig. S13D). Among mossy fibers with lower than 20Hz baseline activity we selected the eye-modulated neurons by filtering out units with less than 70 Hz of change in their response aligned to deceleration onset of targeted saccades. Among 393 remaining mossy fibers, we extracted features to separate goal and state groups. These features included (1) normalized distance measures between the responses to reward relevant and irrelevant saccades while controlling kinematics (direction, amplitude, peak velocity), (2) visual response of the cells 20-50 ms after cue presentation (Fig. S13F), and (3) the shape of amplitude and angle tuning curves (Fig. S13E bottom, Fig. S11E, Fig. 4C). We used UMAP and k-means algorithms to cluster these mossy fibers into two groups (state and goal) using aforementioned features (Fig. S13G).

*Forming cliques based on neuron-neuron spike interactions*

After acquiring the cross-correlograms for each pair of neurons, we jitter-corrected the result (*7*). The result clustered the neurons into small networks, called cliques. The procedure began by forming an adjacency matrix (Fig. 1C), where each neuron was a row and a column in the matrix. We then made the adjacency matrix symmetric by keeping the maximum absolute value of the cross correlogram during the 0-3 ms period for each cell pair (causal window). Next, we used spectral clustering to divide the adjacency matrix into groups. Spectral clustering is a graph clustering method which uses the adjacency matrix of the graph and represents the nodes (each cell) in a 2D or 3D subspace of eigenvectors of the Laplacian matrix (spectral space). In the spectral space the interconnected nodes have shorter distance from each other than the weakly connected nodes (*48*). To illustrate clique membership, we used the weights to assign line thickness and then connected the cell's location on the probe to its clique (Fig. 1C, left subplot). We further verified our cell type identification by ordering neurons in each clique in the adjacency matrix by hierarchical clustering. This algorithm organized the rows of the matrix by putting similar rows next to each other. The result was block sub-matrices of the same cell types with similar interactions within and between cell types.

*Computing the potent vector*

In this region of the cerebellum, the climbing fibers report to the cerebellum the direction of visuomotor events (*10*, *49*), likely because of the superior colliculus projections to the inferior olive (*50*). For example, the climbing fibers reported to the cerebellum both the direction of the visual event, and then independently, the direction of the planned saccade (*14*). We used the properties of the climbing fiber input that the P-cells of a clique received to assign a potent vector to all the neurons that were a member of the clique. The strength of this encoding differed among the various P-cells. We quantified this encoding by measuring the CS rate as a function of the direction of the visuomotor event during two different windows: 40 to 85 ms with respect to the visual cue onset, and -70 to +30 ms window from saccade onset (Fig. 2A, Fig. S5A). We then fitted a Von Mises distribution to the resulting rates as a function of angle (*51*). This produced two vectors, one that quantified the strength and direction of the CS response to the visual events, and a second that quantified the strength and direction of the CS response to the motor events. Each vector had an angle $\theta$ and amplitude $\rho$, where the amplitude identified the sharpness of tuning. Thus, the amplitude of the vector was bounded by 0 (not tuned), and 1 (maximum sharpness). This produced two distinct vectors for each P-cell: one describing the tuning with respect to visual events, and one describing the tuning with respect to motor events (Fig. S5B, left). However, encoding of these two vectors were very similar: the magnitude of the vector for the visual event was highly correlated with the magnitude of the vector for the motor event (Fig. S5B, right). Hence, for each P-cell, we used the vector with the larger amplitude and labeled it as its *potent vector*.

*Testing the validity of the potent vector*

The potent vector was defined based on the climbing fiber information that each P-cell received. However, we conjectured that the direction and magnitude of the potent vector were predictors of the downstream influence that the P-cells had on the eye movements. To examine this conjecture, we performed three tests.

First, we tested our estimate of the direction of the potent vector $\theta$. To do this, we quantified the effects that a CS-triggered SS suppression had on the trajectory of a saccade (Fig. 2B). We observed that the downstream effects, on average, were focused on the direction of the potent vector, with little or no contributions in the direction perpendicular to this vector (Fig. 2C). Moreover, when two P-cells with potent vectors that were nearly parallel were simultaneously suppressed, the effects in the direction of the potent vector roughly tripled in direction $\theta$ with little or no contribution in direction $\theta + \frac{\pi}{2}$ (Fig. 2C). These results implied that our estimate of the direction of the potent vector for each P-cell was an unbiased estimate of the actual direction of the weight vector $\boldsymbol{w}$ that functionally defined the connection between that P-cell and the eye muscles.

Second, we tested our estimate of the magnitude of the potent vector. To do this, we divided our population of P-cells into 3 groups based on the amplitude $\rho$ [edges: $\rho = 0, 0.15, 0.3$]. We observed that as $\rho$ increased between the groups, so did the downstream effects of SS suppression (Fig. 2D). Next, we considered the effects of simultaneous suppression of two P-cells, finding that the size of the displacement produced grew linearly with the size of the amplitude of the sum of the two potent vectors (Fig. S7C). These results implied that our estimate of the amplitude of the potent vector for each P-cell was correlated with the actual amplitude of the weight vector that functionally defined the connection between that P-cell and the eye muscles.

Third, we asked whether simultaneous suppression of two P-cells during a single saccade produced a displacement of the eyes in a direction and amplitude that was predicted by the superposition of the two potent vectors of the P-cells. We observed results that were consistent with this prediction (Fig. 2E).

*Assigning a potent vector to each clique*

We organized the P-cells and the neighboring MLIs into neuronal cliques and then assigned a single potent vector to all the neurons in that clique. To do this, we considered cliques that had at least one P-cell (both SS and CS). We observed that the climbing fibers that projected to the same clique carried information that was much more similar to each other than between cliques (Fig. S5C). For example, the visuomotor tuning distance, measured as the difference in the strength of tuning $\rho$ of visual vs. motor events (distance of two points in Fig. S5B, as shown in Fig. S5C), was much less within a clique than between cliques (rank sum test, p=1.5e-26). Similarly, the difference in the angle $\theta$ of the potent vectors was much less within a clique than between (rank sum test, p=1.12e-18). To compute the potent vector for a given clique, we performed a vector average of the potent vectors of the P-cells that belonged to that clique, then assigned that single potent vector to all the P-cells and interneurons in that clique. Individual climbing fiber's preferred direction differences from the clique's potent direction was centered at zero, as was the difference in the amplitude (Fig. S5D-E).

*Perturbation of saccades following a CS-triggered SS suppression*

For this analysis we used the spike-triggered averaging techniques introduced in (*17*). We focused on the saccades to the primary target as they had a fixed amplitude and started from the center fixation point. For each P-cell, we divided saccades that were made from the same initial position to the same target into two groups: (1) saccades in which the P-cell did not experience a CS during the time window -60 to +40ms with respect to saccade onset (NOCS: no cs), (2) saccades in which the P-cell experienced a single CS during the time window -30 to +30ms with respect to saccade onset (WCS: with cs). Then, we took steps to ensure that the start positions of the saccades were comparable between WCS and NOCS saccades. For each target direction, we computed the initial position of each WCS and NOCS saccade with respect to the fixation point and projected this vector onto the potent vector. We removed the minimum number of NOCS saccades required to match their mean projected start position to within ±0.03 degrees of the mean projected start position of the WCS saccades. We then computed the difference in saccade velocity vectors in each direction as a function of time between WCS and NOCS saccades, projected onto the potent vector, and the orthogonal vector (null vector). For each target direction, this produced two scalar quantities, i.e., change in velocity, as a function of time. We then averaged the projected changes in velocity across all saccade directions. The null distribution was formed through bootstrapping of the populations of P-cells with randomized tagging of trials, labeling them as WCS saccades or NOCS saccades (number of bootstraps=1000).

In order to confirm that the difference in velocity between the WCS and NOCS groups was not due to planning, we verified that the CS-triggered SS suppression displaced the eyes only if the suppression event was within a short window centered on the saccade itself. Indeed, the SS suppression events that occurred longer than 50 ms before saccade onset had no effects on the saccade (Fig. S7A).

*Testing superposition of the potent vectors*

Neuropixels allowed us to record from multiple folia simultaneously, producing cliques that had diverse potent vectors. For example, in Fig. 1C, clique 1 has a large amplitude potent vector, whereas clique 5 has a very small potent vector. Moreover, the direction of the potent vector for clique 1 is near 135$^{o}$, whereas for clique 3 it is near 180$^{o}$. Because the climbing fiber inputs to all of these cliques were recorded simultaneously, we performed spike-triggered averaging on simultaneous complex spikes. This allowed us to ask whether simultaneous suppression of two P-cells produced a displacement that obeyed superposition in the vector space of the two potent vectors.

For two simultaneously recorded P-cells, we had two potent vectors, with properties $(\rho_1, \theta_1)$ and $(\rho_2, \theta_2)$. Next, we selected pairs of P-cells in which the size of the two potent vectors exceeded a minimum threshold: $\frac{\rho_2}{\rho_1} \geq 0.5$, where $\rho_1 > \rho_2$. This ensured that the P-cell with the small potent vector $\rho_2$ would not be completely overwhelmed by the P-cell with the larger potent vector $\rho_1$. We then divided the saccades that were made from the same initial position to the same target into two groups: (1) saccades during which neither of the two P-cells experienced a CS (NOCS, -60 to +40ms window with respect to saccade onset), (2) saccade during which both of the P-cells experienced a single CS within 30ms of each other (WCS, -30 to 30ms window). As in the case of single P-cells, we ensured that the starting positions of the two saccades were matched (within ±0.03 degrees). We then tested the hypothesis that the CS-triggered change in saccade trajectory was the vector summation of the two potent vectors, i.e., $\rho_{net}\angle\theta_{net} = \rho_1\angle\theta_1 + \rho_2\angle\theta_2$. The null distribution was formed through bootstrapping of the populations of P-cell pairs with randomized tagging of trials, labeling them as WCS saccades or with NOCS saccades (number of bootstraps=1000).

*Computing a potent vector for neurons in a clique*

For clique j, we computed the potent vector as the average vector defined by its climbing fiber inputs (only P-cells with both SS and CS):

$$\rho_j\angle\theta_j = \frac{1}{n_j} \sum_{i\in j} \rho_i\angle\theta_i \tag{1}$$

In the above equation, $n_j$ is the number of P-cells in the j-th clique. For each of the cell types in a clique we first binned the saccades based on the movement angle to 8 directions (0:45:360 deg). We then averaged CS responses to saccade directions with respect to $\theta$ (Fig. 3A-B). Instantaneous firing rates were calculated from peri-event time histograms with 1 ms bin size. We used a Savitzky–Golay filter (2nd order, 31 datapoints) to smooth the traces for visualization purposes.

*Computing population responses*

When a P-cell is suppressed, its downstream effect on behavior is specified by its potent vector, i.e., a force in direction $\theta_i$ with a magnitude $\rho_i$. For the eyes, the premotor neurons are the horizontal and vertical burst generators. P-cells project to these neurons indirectly; we represent the downstream effects of a P-cell on the horizontal and vertical burst generators $(F_x, F_y)$ using weights $w_x, w_y$. When a saccade is made in direction $\psi$, a P-cell fires with the rate $r_i(\psi, t)$ : that neuron's contribution to that saccade is:

$$F^i = \left[F_x^i, F_y^i\right] = -\left[w_x, w_y\right]^T r_i(\psi, t) = -\frac{\rho_i}{\sum_j \rho_j} [\cos(\theta_i), \sin(\theta_i)]^T r_i(\psi, t) \tag{2}$$

Because the direction $\theta_i$ (referring to the potent direction of the P-cell) depends on the anatomical location of each neuron in the cerebellum, we must remove our sampling bias. To explain this, note that cells on the right of the midline tend to have a potent vector toward the left, and cells on the left of the midline tend to have a potent vector toward the right (*11*). This implies that there is a need to account for the anatomical position dependence of the neurophysiological data.

To account for this anatomical variability, we assumed that the true distribution of the potent vectors uniformly covered the direction space. That is, we assumed that for each P-cell that we had recorded, there were 7 others that were identical in every way except one critical property: their potent vector was $\frac{\pi}{4}, \frac{\pi}{2}, \frac{3\pi}{2}$, ... away from the current cell. Thus, we duplicated each neuron $i$ by uniformly distributing $\theta_i$ as 8 P-cells:

$$\tilde{F}^i = -\frac{\rho_i}{\sum_j \rho_j} \sum_{n=0}^{7} \left[\cos\left(\theta_i + \frac{n\pi}{4}\right), \sin\left(\theta_i + \frac{n\pi}{4}\right)\right]^T r_i\left(\psi + \frac{n\pi}{4}, t\right) \tag{3}$$

The resulting force of population, $P$, is simply the sum of the pseudo-population response from each individual neuron: $P = \sum_i \tilde{F}^i$.

It is noteworthy that, without loss of generality, we can rotate the space for each neuron so that the saccade direction is represented with respect to the direction of its potent vector $\theta_i$. This is equivalent to a simple change of variable $n = \tilde{n} - \frac{\theta_i}{\frac{\pi}{4}}$ ($\theta_i$ is one of the 8 directions). Hence the population response $P$ can be rewritten as follows:

$$P = \sum_i \tilde{F}^i = \sum_i -\frac{\rho_i}{\sum_j \rho_j} \sum_{n=0}^{7} \left[\cos\left(\frac{n\pi}{4}\right), \sin\left(\frac{n\pi}{4}\right)\right]^T r_i\left(\tilde{\psi}_i + \frac{n\pi}{4}, t\right) \tag{4}$$

Where $\tilde{\psi}_i = \psi - \theta_i$ is the saccade direction aligned to the potent vector of the neuron. Hence, assuming isotropic responses and given downstream weights $(w_x, w_y)$ for each P-cell, the net force generated by the population ($P$) can be represented as a weighted projection of the rates $(r_i(\tilde{\psi}_i, t))$ on the $\theta$ and $\theta + \frac{pi}{2}$ axes. Hence, forming the population of P-cells by rotating saccade direction with respect to $\theta$, i.e., the potent vector or CS-on direction of the P-cell, as suggested in (*14*), is equivalent to assuming isotropy for each observed P-cell (Eq. 3).

Figs. 3A & 3B show the average responses of each cell type aligned to the direction of the potent vector $\theta$, i.e. $\bar{r}(\tilde{\psi}_i, t)$. The interesting fact is that while the potent vector for each P-cell is along direction $\theta$, the cells are active in all directions. This implies that when a saccade is toward direction $\psi$, there are many P-cells that are firing and whose downstream forces are pulling the eyes not along $\psi$, but along $\psi + \frac{\pi}{2}$. If the null space theory is right, the spikes in these neurons must be fully canceled. Indeed, both bursters and pauser P-cells in our dataset show a symmetrical response around direction $\theta$, with the firing rates for direction $\theta + \frac{\pi}{2}$ and $\theta - \frac{\pi}{2}$ cancelling each other (Fig. 3B). In addition, the responses towards directions $\theta$ and $\theta + \pi$ show a differential pattern that is time locked to the saccade offset.

As a result, in the population response for the P-cells, the rates parallel to the direction of the movement exhibited a burst-pause pattern that crossed zero at deceleration onset, while in the direction perpendicular to the movement nearly all the spikes canceled. We calculated the confidence intervals for this zero-crossing using bootstrapping. To do so, we calculated the first zero-crossing in a -30ms to 75ms window around deceleration onset for bootstrapped population (100 times with replacement). We repeated the same procedure for finding MLI1 population peak time in a -10ms to 50ms window (Fig. 4I, S14C). In Fig. S19B, we performed the same analysis by shifting the search windows by 15ms and -25ms for onset and offset respectively.

*Estimating the duration of modulation for individual neurons vs. the population response*

Neurons in the population produced cancellation of spikes so that the duration of modulation was precisely aligned to movement duration. To quantify this and compare it to modulation in the individual neurons, we first grouped the P-cells into bursters, pausers, and combined. In addition, we considered the population response $p_x(t)$ (Fig. S19A, similar to Fig. 3A). For each group we found the onset and offset of the average rate modulation as a function of direction with respect to the potent vector $\theta$. Onset and offset times were found by calculating the time points in which the absolute derivative changed in time (MATLAB *findchangepts* function). For each group we bootstrapped the cells to estimate the distribution of onset and offset times in the -100ms to 150ms window around deceleration onset (Fig. S17, n=1,000). The combination of bursters and pausers showed an earlier offset closer to saccade end in comparison to divided groups of pausers and busters in the ipsilateral direction. In addition, the projected population $p_x(t)$ had closer modulation end to saccade offset (mean±SEM: bursters: 88.6±0.17ms, pausers: 77.6±0.23ms, all: 50.1±0.73ms,: $p_x(t)$ 30.8±0.29ms from deceleration onset, ANOVA on modulation offset in +180° for n=1,000 bootstraps per group, F(3)=3904, p<1e-100; saccade offset: 23.3±0.01ms).

*Estimating potent vectors using principal component regression*

We next asked if the downstream weights can be estimated using the original approach used in Kaufmann et al. (*2*) (see Fig. S1). This approach aims to find a linear map from neural activity $N$ on to behavior or muscle activity $M$. For muscle activity, instead of using EMG, we used horizontal and vertical kinematics of the eye (acceleration and velocity). This assumption is quite reasonable as a proxy for premotor and motorneuron control of eye movements (*52*). In order to prevent recording location bias in the P-cells, following aforementioned isotropy assumption, we formed neural activity matrix $N$ by adding 7 rotated copies of each P-cell in space. The matrix $N$ is of size $n \times ct$, where n is number of P-cells times 8 (8 rotations of each P-cell's response), c is number of directions of saccades, and t is the number of time points aligned to saccade maximum velocity (Fig. S21A). We then found the first 10 principal components of the neural activity, labeled as $\Phi$. The first and second principal components represent burst/pause activity irrelevant of the variability of response to different directions (*42*).

We then used ridge regression with cross-validation to find the projected weight matrix $W$ that mapped the neural activity either to velocity, or acceleration. We further used the weight matrix to find the contribution of each cell to the x and y direction of the movement. Converting these contributions to the polar coordinate (angle and amplitude) was significantly correlated with individual cell's potent vector defined based on the Complex Spike activity only if acceleration was used as matrix M. The results were not correlated with the hypothesized potent vector in case of fitting the velocity of the eye.

To summarize, if we assumed that the $M$ matrix was eye velocity, which is consistent with activities of motoneurons, then the weight vector for each neuron that we found through regression had no relationship to the potent vector that we found using their climbing fiber inputs. However, if we assumed that the $M$ matrix was acceleration, then the weight vectors were highly correlated to the potent vectors. Thus, the ability to recover the potent vector depends on having the correct assumption regarding what is being computed by the network.

*Statistical testing*

We performed t-tests, or rank sum tests, or ANOVAs, to compare distributions. To compute effects of SS suppression, we computed the null hypothesis via boot-strapping, randomly assigning labels to the data.

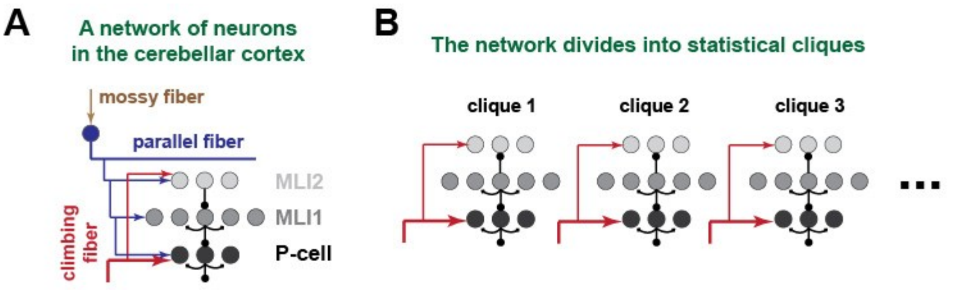


**C**

behavior — potent vectors — measured neural activity (P-cells or MLI)

$$\underset{2 \times T}{P} = \underset{2 \times n}{W}\,\underset{n \times T}{N}$$

| Typical approach | Current approach |
|---|---|
| **1. Measure simple spikes in P-cells and MLIs**<br>**2. Measure behavior**<br>**3. Find W by mapping neural activity onto behavior** | **1. Measure complex spikes in each P-cell and fit to behavior to find W.**<br>**2. Verify that the W computed from complex spikes is an unbiased predictor of the change in behavior when the simple spikes of a P-cell are suppressed.**<br>**3. Verfiy that superposition holds in behavioral space as predicted by W when the simple spikes of two P-cells are simultaneously suppressed.**<br>**4. Using the W from the complex spikes, project the simple spikes of the P-cells onto an output space.**<br>**5. Find MLIs that belong to the same "clique" as a group of P-cells and have the MLIs inherit the potent vector associated with those P-cells.**<br>**6. Using the inherited W, map the MLI activity onto the output space.** |

**Fig. S1**. **Outline of the theoretical approach. Our objective is to understand what is being computed by a network of neurons in a region of the cerebellum concerned with eye movements. (A)** The network is composed of a 3-layer system of neurons in which mossy fibers and climbing fibers provide the inputs onto two groups of interneurons (MLI1s and MLI2s) and one group of output neurons (P-cells). MLI2s inhibit MLI1s, and MLI1s inhibit the P-cells. The P-cells produce their output via simple spikes at firing rates of 50-100 Hz. The climbing fiber input is a stochastic, low frequency (1 Hz) signal that briefly but completely suppresses the simple spikes. **(B)** Using spike interactions among simultaneously recorded neurons, the network separates into cliques, i.e., small clusters of tightly coupled neurons. **(C)** The typical approach measures spiking in the P-cells and the MLIs and then maps them onto behavior, e.g., eye velocity, or acceleration, in Cartesian space. The result of this fitting is the matrix $W$ which for each cell contains a 2-dimensional (2x1) vector $\boldsymbol{w} = [\boldsymbol{w}_x, \boldsymbol{w}_y]$, termed the potent vector. Here, we first measure the climbing fiber input to each clique, i.e., the 1 Hz complex spikes, and fit it to behavior, finding a vector $\boldsymbol{w}$ for each P-cell. Using spike-triggered averaging on the climbing fiber input, we verify that when a P-cell's simple spike production is briefly suppressed, the downstream effect on behavior is, on average, a displacement of the eyes as predicted by $\boldsymbol{w}$**.** Then, we verify that when two P-cells receive simultaneous climbing fiber inputs, the resulting simultaneous simple spike suppression moves the eyes as predicted by the superposition of their two $\boldsymbol{w}$ vectors. Next, we use the climbing fiber estimate of $\boldsymbol{w}$ to form the $W$ matrix and now map the P-cell simple spikes onto an output matrix $P$. This output is interpretable, illustrating a signal that specifies when and how to stop the ongoing movement. Critically, the simple spikes that would move the eyes away from the intended target are present in the individual P-cells but canceled in the population. Finally, we assume that all the interneurons that are in a clique inherit the $\boldsymbol{w}$ vector of the P-cells in the same clique. We map the activities of the interneurons using the clique's $\boldsymbol{w}$ vector, finding an interpretable outcome that transforms the mossy fiber inputs to the P-cell output.

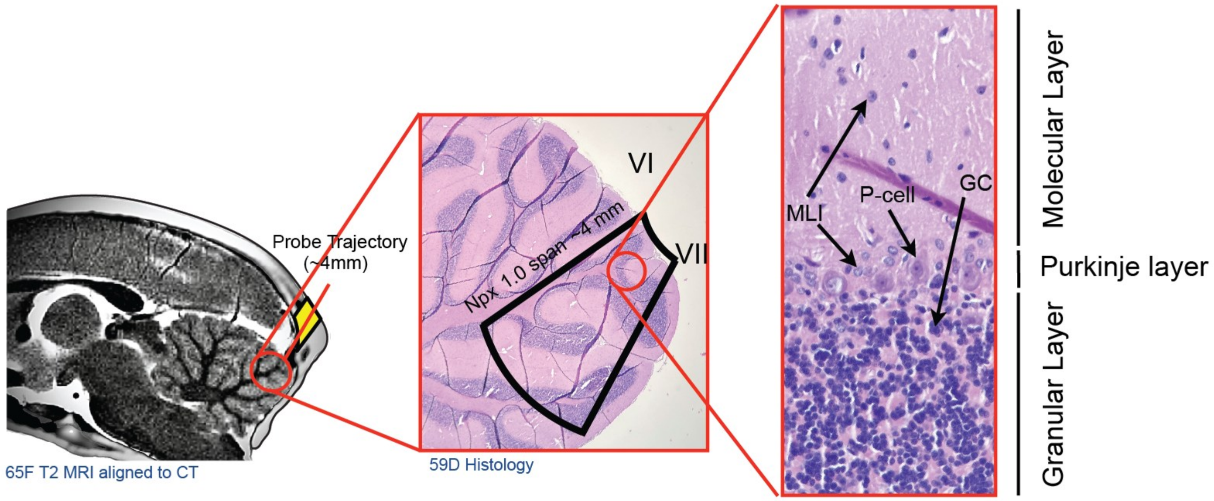


**Fig. S2**. **Electrophysiology paradigm and recording regions.** (left) T2 MRI image overlaid by CT scan of Monkey B (midline section). (middle) Histology section of Monkey R (59D) with approximate probe trajectories and span of recording. (right) histology image of 3 layered structure of cerebellar cortex.

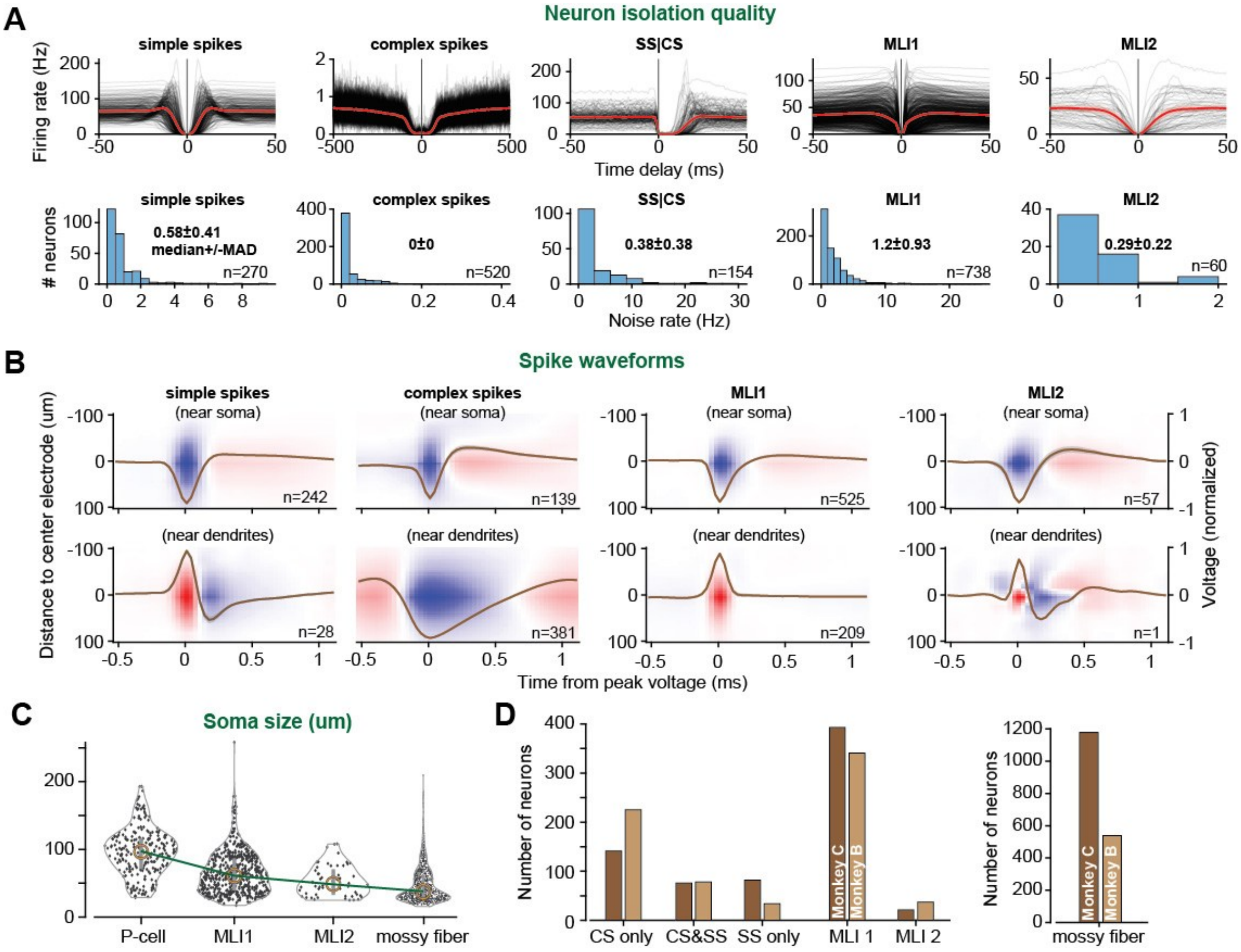


**Fig. S3**. **Isolation quality of the neurons in the data set. (A)** Top: Auto-correlograms (and SS|CS cross correlogram) of each cell type with average traces (red). SS|CS describes the conditional probability of an SS, given that a CS occurred at time zero. The probabilities are for bins of 1 ms duration and are multiplied by 1000 to represent spike rate in Hz. Bottom: refractory violation distribution of for each cell type (<1ms for all neurons except complex spikes and SS|CS: <5ms). **(B)** Normalized spatiotemporal waveforms of each cell type plotted as a function of distance of the electrode with respect to the electrode that recorded the largest waveform (center electrode). The waveforms are divided into somatic (downward) and dendritic (upward) waveforms. Brown lines indicate the maximum waveform (distance = 0 um from the best electrode). **(C)** Cell size estimates using FWHM of the spatiotemporal waveforms at time = 0 ms for somatic spikes. **(D)** Number of each cell type recorded in each monkey. Error bars are 2xSEM.

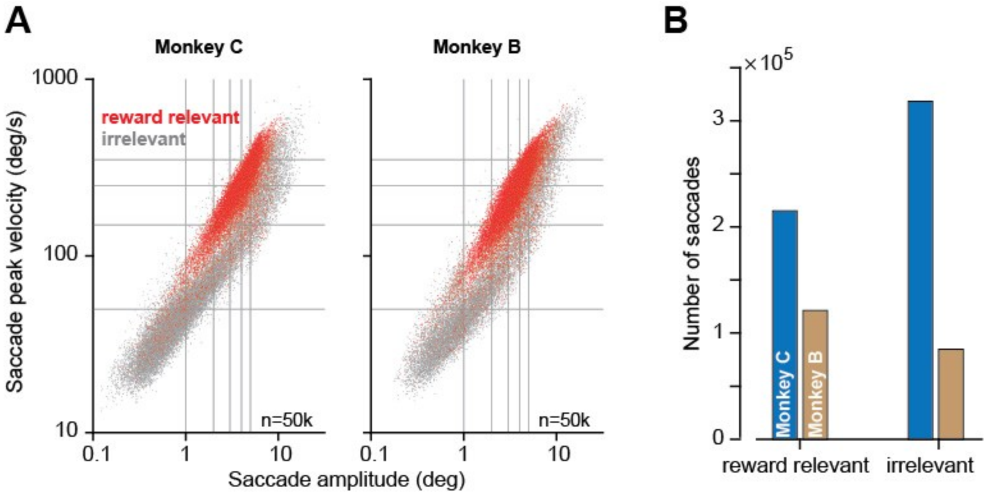


**Fig. S4**. **Kinematic properties of saccades of the two monkeys in the data set. (A)** Main sequence plots in log-log scale. Red dots indicate reward relevant saccades (primary, secondary and back to center saccades, see Fig. 1A). Gray dots indicate reward irrelevant saccades. **(B)** Number of reward-relevant and irrelevant saccades for each subject. The data in part A were randomly sampled from the roughly 0.7M saccade database shown in part B.

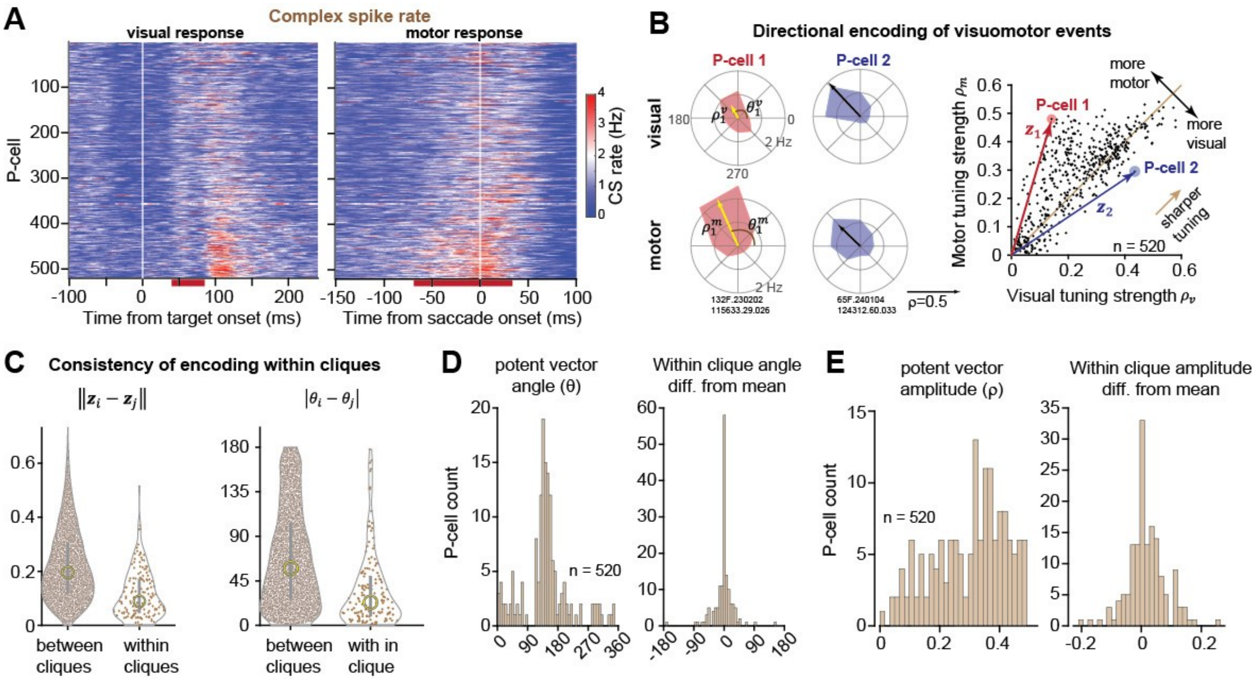


**Fig. S5**. **The climbing fiber input to a P-cell defined a vector space. This input was consistent for neurons within a clique. (A)** Complex spike response in the potent direction sorted by motor minus visual response, aligned to target onset (left) and saccade onset (right). The red lines at the bottom indicate the period for which the visual and motor responses were measured (40-85ms after target onset, -70 to 30 ms with respect to saccade onset). **(B)** The CS rates were measured as a function of direction of the visual and motor events and each fitted to a von Mises distribution. The result for each P-cell is a vector with amplitude $\rho$ and direction $\theta$. For each P-cell, a separate vector was found for the visual and motor responses, $\rho^v \angle \theta^v$ and $\rho^m \angle \theta^m$. In the two neurons on the left subfigure, the potent vectors are indicated, demonstrating that in neuron 1 the climbing fiber input had a stronger motor response, whereas in neuron 2 the visual and motor responses were similar. On the right subplot, the vector $\boldsymbol{z} = [\rho^v, \rho^m]$ indicates the amplitudes of the visual and motor potent vectors for each P-cell. The results show that while visual and motor responses were highly correlated, most P-cells received stronger information from the olive regarding movements rather than visual events. **(C)** (left) Euclidean distance between $\boldsymbol{z_i}$ and $\boldsymbol{z}_j$ (as shown in the right panel of par B), for two P-cells within and between cliques. (right) For each P-cell we defined its potent vector to be the larger of the visual and motor potent vectors. Thus, each P-cell was assigned a vector $\boldsymbol{w}$ with amplitude $\rho$ and angle $\theta$. Right plot shows the absolute angle difference between the potent vectors within and between cliques. The P-cells within a clique received more similar information from the inferior olive than cells between cliques. **(D)** (left) Distribution of $\theta$. The distribution is not uniform, which indicates that there is an anatomical bias in the recordings. (right) Distribution of angle difference in the potent vectors of each clique with respect to the mean angle. **(E)** (left) Distribution of the amplitude of the potent vectors $\rho$. (right) Distribution of magnitude difference of the potent vectors within a clique with respect to the clique's average $\rho$.

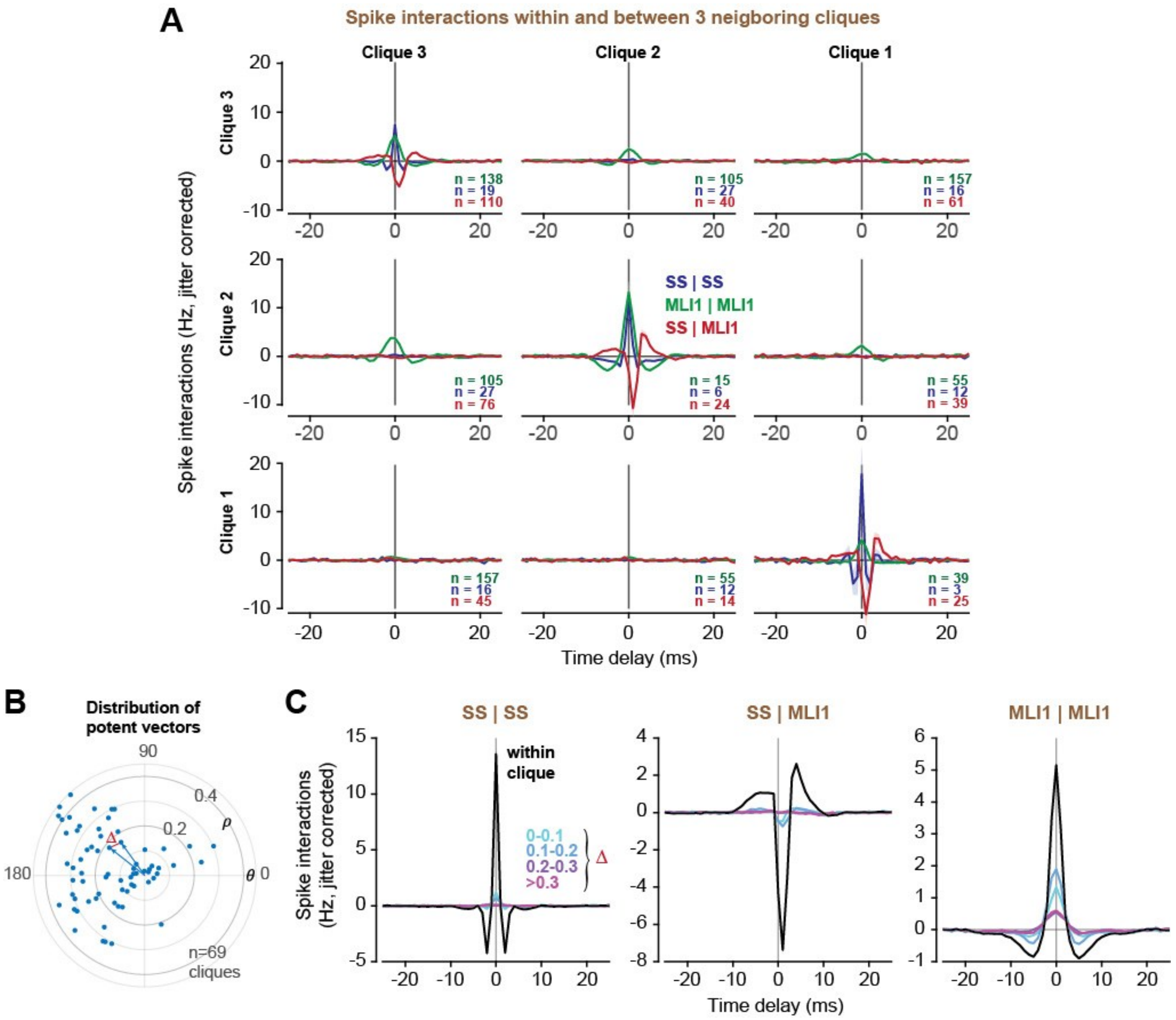


**Fig. S6. Cliques defined a finer unit of computation in the cerebellum than microzones**. Microzones are defined based on the similarity of the information that they receive in their climbing fiber inputs (*15*). Here, we quantified how spike interactions among neurons varied between cliques as a function of the difference in their climbing fiber inputs. **(A)** Data from a single recording (Fig. 1), focusing on cliques 1-3 which had similar climbing fiber inputs. The matrix shows interactions between neurons of each clique. The diagonal plots are within clique interactions. For example, SS|SS is the probability of synchronous simple spikes in two P-cells, at various delays, corrected for chance via jittering. SS synchrony is present only when the P-cells are in the same clique. **(B)** The distribution of potent vectors for the cliques in the data set. The potent vectors are derived from the information in the climbing fiber inputs to each clique. Each vector has an amplitude $\rho$ and direction $\theta$. The distance between two vectors is specified by $\Delta$. **(C)** Spike interaction between cliques as a function of difference in their potent vectors. For example, SS synchrony drops by an order of magnitude when two P-cells are not in the same clique, despite have very similar climbing fiber input (as measured by $\Delta$). Data in this subplot includes only the recordings in which multiple P-cells and MLIs in more than one clique were simultaneously isolated (n=44 cliques).

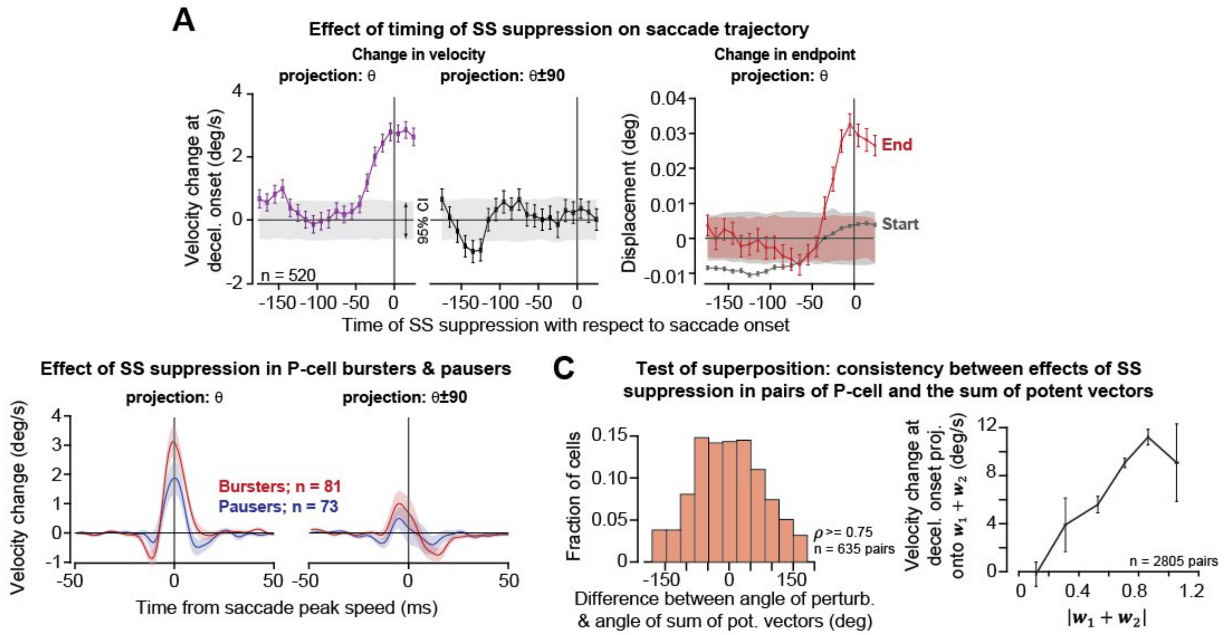


**Fig. S7**. **The potent vector for a P-cell was an unbiased estimator of the magnitude and direction of the downstream effects that SS suppression had on behavior. (A)** CS-triggered SS suppression perturbed the eyes only if it took place near saccade onset or during the saccade. Left subplot shows the effect on velocity of the saccade as a function of when the P-cell was suppressed with respect to onset of the saccade (30ms moving window). The vector of velocity change was measured for each P-cell $i$ at peak saccade velocity, then projected onto $\theta_i$ (direction of the potent vector), and $\theta_i + \frac{\pi}{2}$. Right panel shows the effect of SS suppression on the start and end position of the saccade as a function of when the P-cell was suppressed with respect to saccade onset. SS suppression was followed with an overshoot of the saccade in direction of the potent vector. Displacement and velocity changes are with respect to movements in which the P-cell was not suppressed. **(B)** The effects on saccade trajectory when the P-cell was suppressed, separated for burster type and pauser type P-cells (WCS: -30 to 30ms with respect to saccade onset, NOCS: -60 to 40ms). In both cases, P-cell suppression produced a pull in direction of its potent vector. **(C)** Test of superposition: the potent vector was an unbiased estimate of the effects of SS suppression on behavior. This analysis relied on pairs of P-cells that were suppressed within 30 ms of each other (WCS: -30 to 30ms, NOCS: -60 to 40ms, with respect to saccade onset). Left subfigure shows the difference between the angle of the vector of velocity change caused by SS suppression (measured at saccade peak velocity), and the angle predicted by the sum of the two potent vectors (mean±SEM, -0.474±3.16). We picked P-cell pairs for which the angle of the potent vectors was within $\theta_2 - \theta_1 \leq \frac{\pi}{4}$, and $\rho_1 + \rho_2 \geq 0.75$. Right subfigure shows that as the amplitude of the sum of the two potent vectors for the pair of P-cells increased, so did the magnitude of the velocity change vector that was caused by simultaneous SS suppression of those two P-cells. Here, the two P-cells have potent vectors that are within $\frac{\pi}{8}$ of each other. Error bars are SEM.

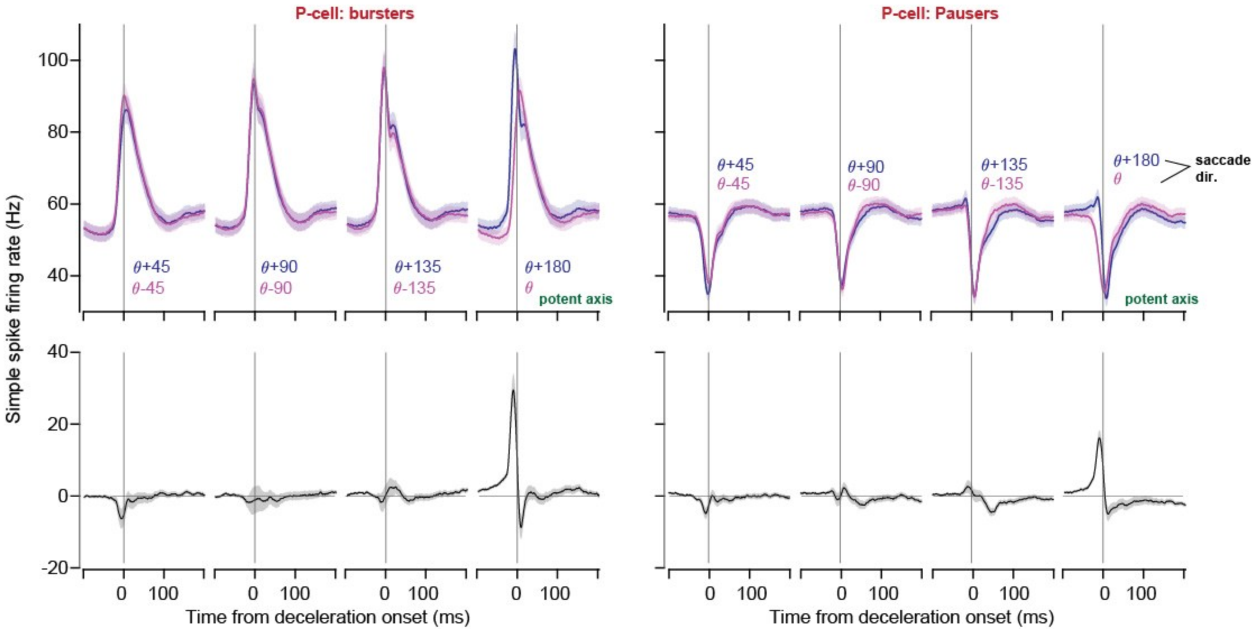


**Fig. S8**. **P-cell bursters (left) and pausers (right) exhibited simple spike symmetry with respect to their potent vector.** For each P-cell, the angle of the potent vector is specified by $\theta$. Saccade direction is with respect to the P-cell's potent vector. (top) Average SS responses of P-cells during saccades with respect to the angle of the cell's potent axis. (middle) Same as top row, but for opposite direction saccades with respect to the potent axis. (bottom) Subtraction of the top and middle responses. In the left and right subplots, the 4th column shows the differential response on the axis of symmetry (potent axis).

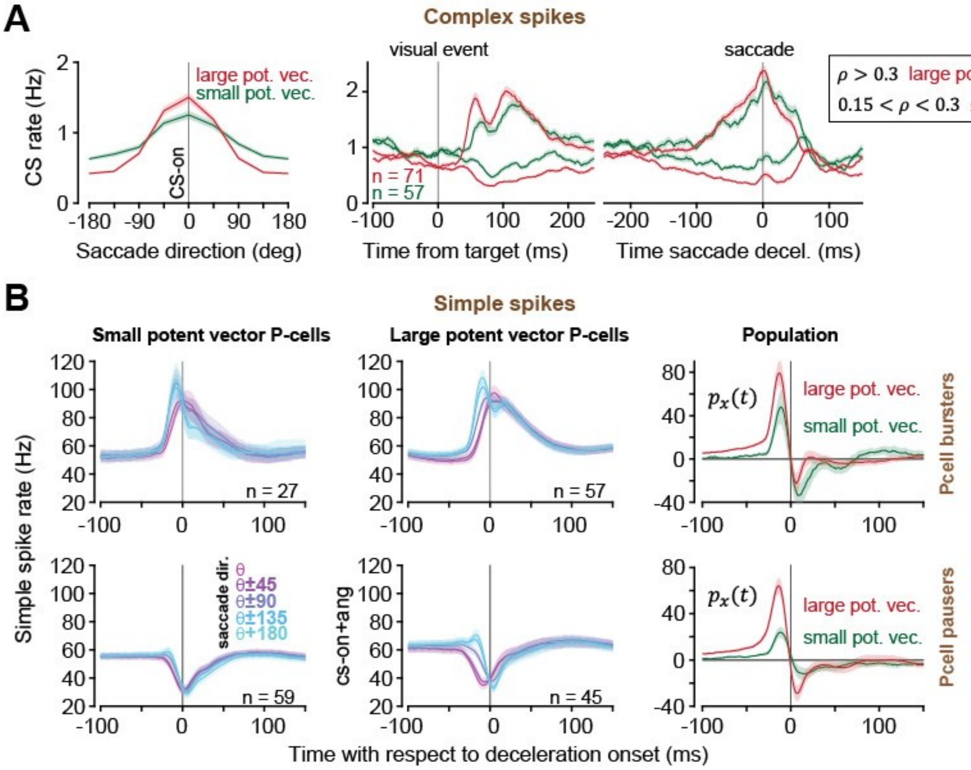


**Fig. S9**. **Cliques with larger potent vectors showed larger population responses, despite having similar SS rate modulations. (A)** (left) tuning curve of the larger ($\rho > 0.3$, red) and smaller ($0.15 < \rho < 0.3$) potent vector P-cells. (right) CS responses of the two groups aligned to target onset (left) and saccade deceleration onset (right) for the potent direction $\theta$, and direction $\theta + \pi$. **(B)** (top row) P-cell simple spike bursters average response for all directions of saccade aligned to the potent axis for small potent vector (left) and large potent vector P-cells (middle column). (right) Population response for two groups in the saccade direction. (bottom row) Same as top row, but for pauser type P-cells. Thus, despite the fact that the P-cells in both groups produced similar modulation of simple spikes, in the population, more of the spikes produced by the small potent vector P-cell group were eliminated.

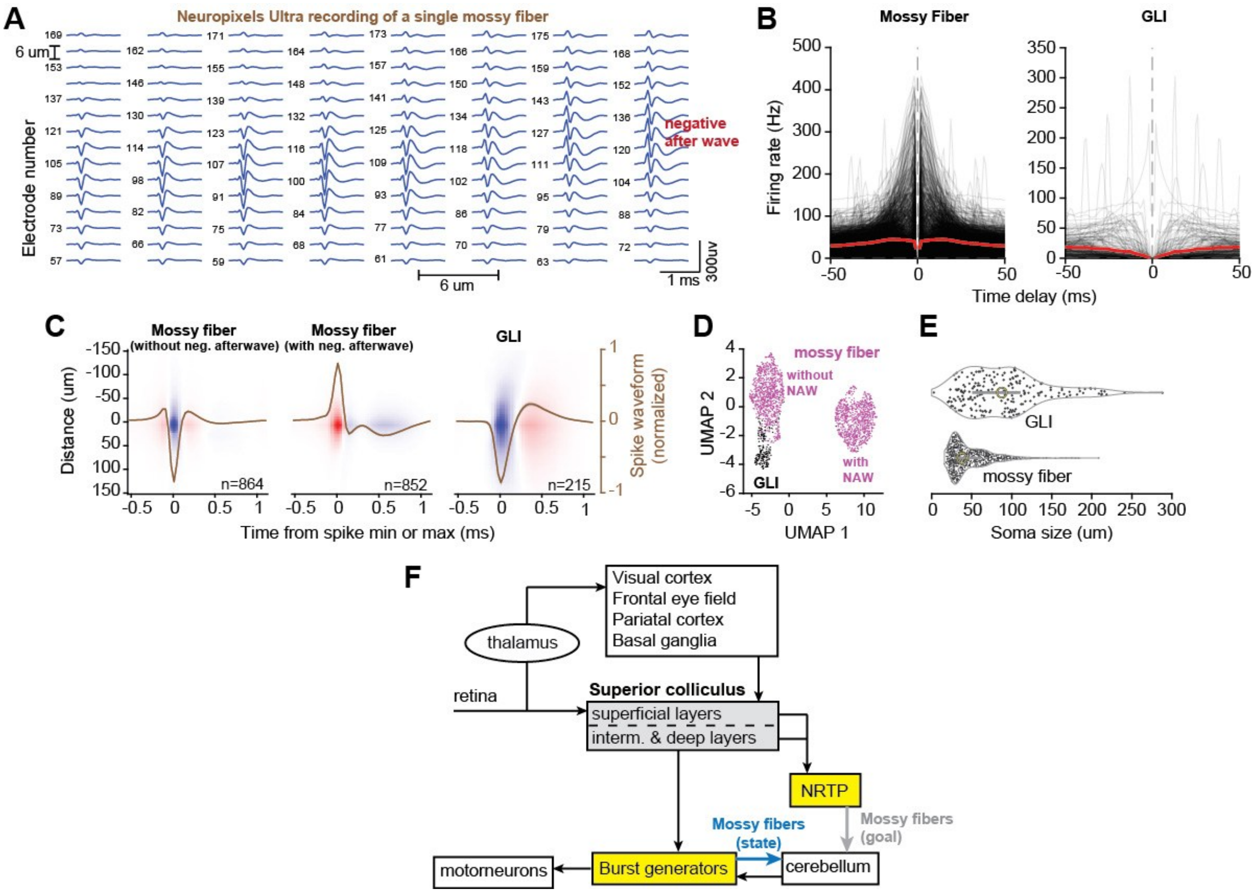


**Fig. S10**. **Mossy fibers were detected in the granular layer by their triphasic waveform with negative after wave.** **(A)** Neuropixels Ultra recording from a single mossy fiber rosette structure, showing the neuron's spike waveforms on the various channels. Note that depending on the electrode's location with respect to the rosette structure, the waveform changed significantly. On the left side of the probe, this mossy fiber had a negative waveform without an after wave. On the right side of the probe, it had a positive initial peak in its waveform then a negative after wave. **(B)** Auto-correlogram plots of mossy fibers and other granular layer interneurons (GLI). **(C)** Average spatiotemporal waveform of mossy fibers (left: downward without a negative after wave, right upward with a negative after wave) and GLIs. Brown line indicates the average waveform at the center of the heatmap (distance = 0 um with respect to the main electrode). **(D)** Spatiotemporal waveforms of the mossy fibers and GLIs in the UMAP space. NAW: negative after wave. **(E)** Estimated soma size of the mossy fibers (downward spikes) and GLIs. **(F)** Schematic of mossy fiber inputs to the oculomotor region of the cerebellar vermis. NRTP: nucleus reticularis tegmenti pontis. The mossy fiber information includes a copy of the motor commands and a copy of the goal location of the movement.

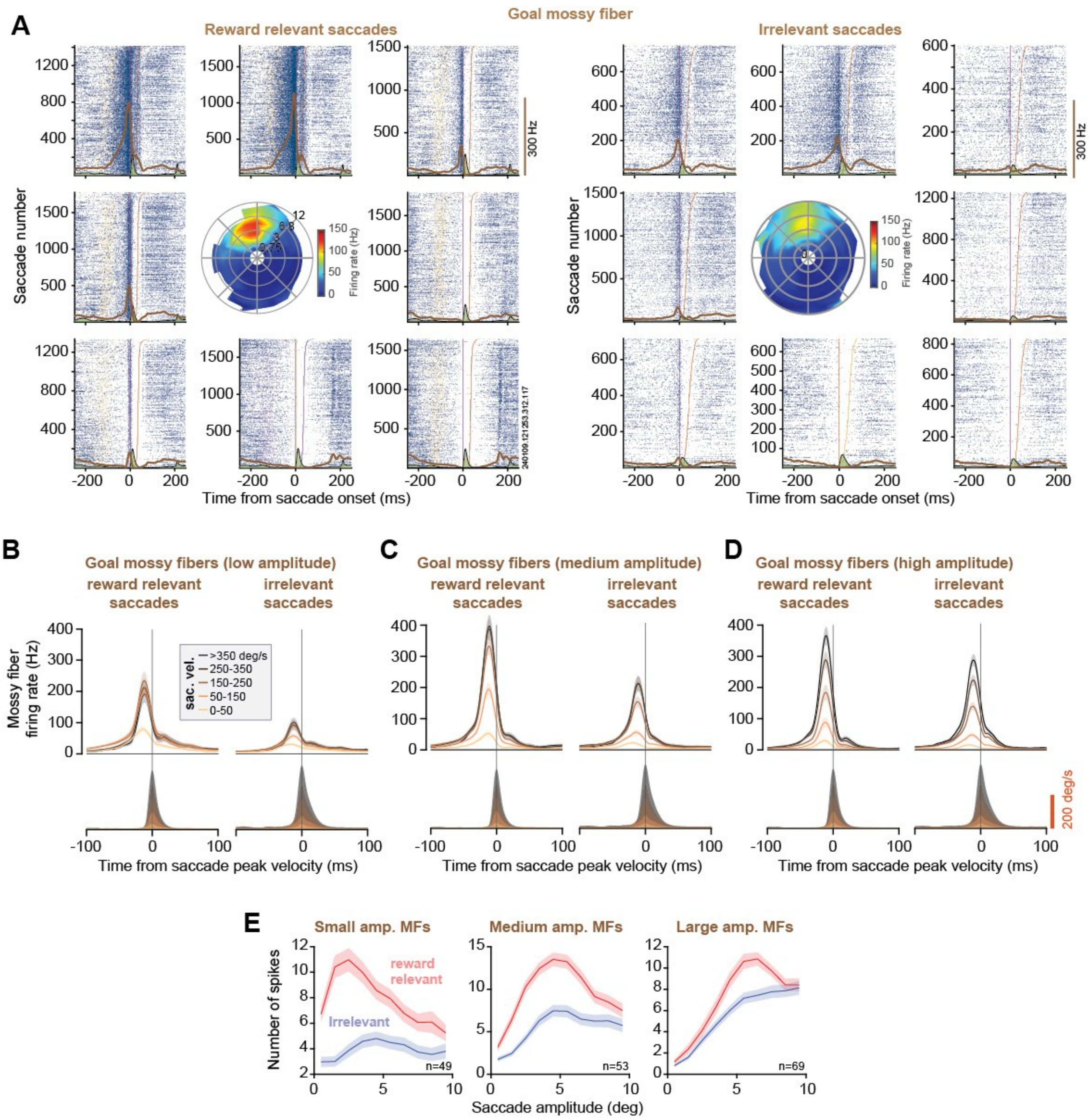


**Fig. S11. Some mossy fibers provided information regarding the goal location of the saccade. (A)** Example of a goal encoding mossy fiber. The responses are shown during reward relevant (left) and irrelevant saccades (right). Each of the 8 rasters represents responses for saccades toward that direction. Center heatmap shows the rate of spikes in the mossy fiber as a function of amplitude (in degrees) and direction of the movement. For reward relevant saccades this mossy fiber had a response field centered at amplitude 4 degrees and direction 100 degrees. For irrelevant saccades the encoding had a similar spatial preference but a much weaker response. **(B)** Mossy fibers that encoded small amplitude goals (data aligned to the preferred direction). The neural responses in the two columns (reward relevant and irrelevant saccades) are for equal amplitude movements. **(C)** Similar to part B, but for the group of mossy fibers that encoded medium amplitude goals. **(D)** Similar to part B, but for the group of mossy fibers that encoded large amplitude goals. **(E)** Number of spikes (-50 to 110ms period with respect to saccade onset) as a function of saccade amplitude for MFs that encode small, medium, or large amplitude goals. Error bars are SEM.

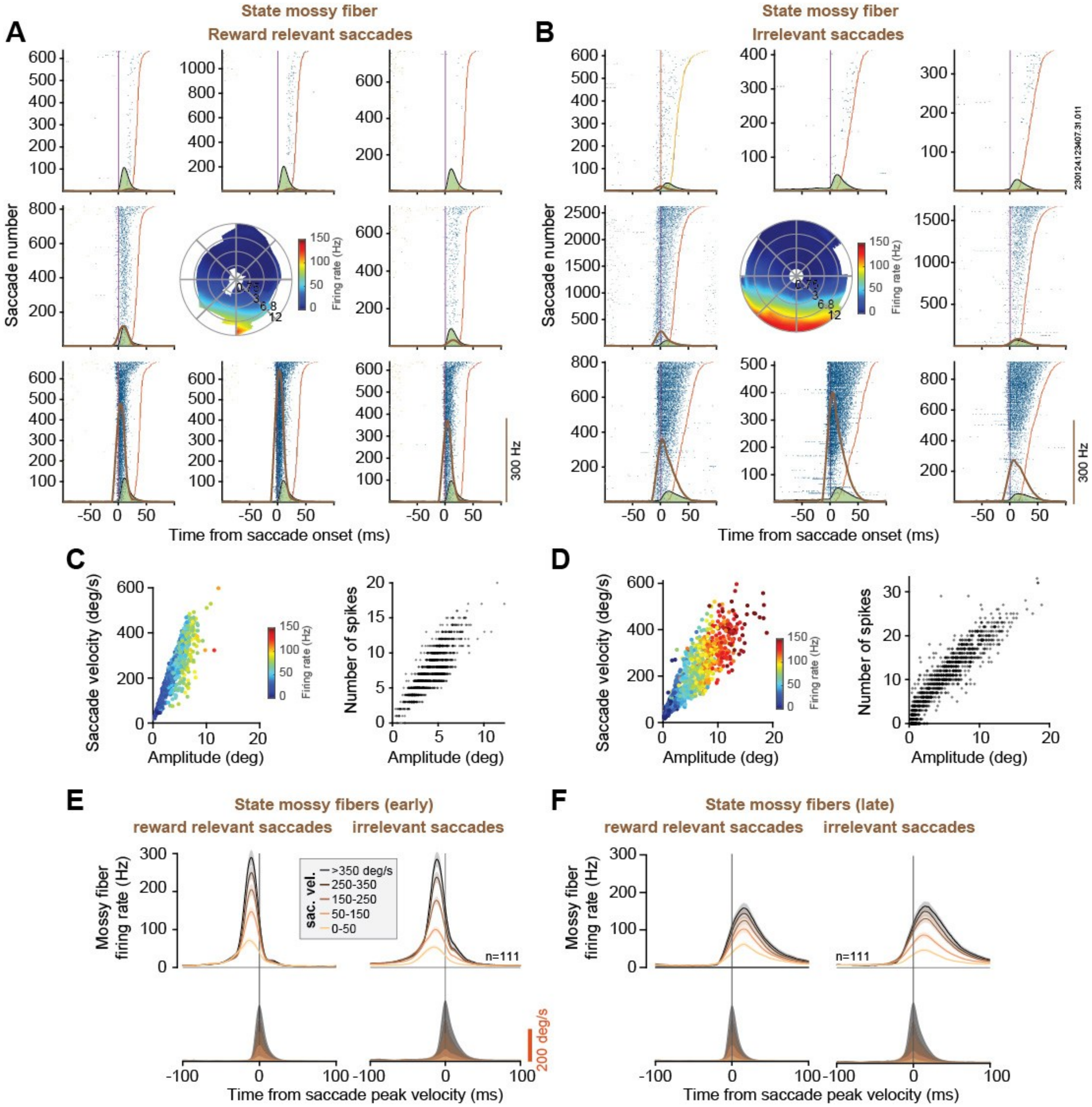


**Fig. S12**. **Some mossy fibers provided information regarding the motor commands. (A) and (B) show examples of a state mossy fiber that encoded eye velocity.** The responses are shown during reward relevant and irrelevant saccades. Each of the 8 rasters represents responses for saccades toward that direction, aligned to saccade onset. Center heatmap shows the rate of spikes as a function of amplitude (in degrees) and direction of the movement. In this mossy fiber, the spike rate grows for downward saccades as a function of saccade amplitude. Notably, the responses are not different for the two types of saccades. **(C)** (left) Velocity-amplitude plot of reward relevant saccades color-coded by the average rate of the mossy fiber for each saccade towards the preferred direction of the mossy fiber. (right) Number of spikes as a function of amplitude of saccades for saccades toward the preferred direction. **(D)** same as (C) for the irrelevant saccades. The integral of the number of spikes is a good estimate of the saccade amplitude for both reward relevant and irrelevant saccades. **(E)** The group of mossy fibers that encoded motor commands with short latency (data aligned to the preferred direction). The neural responses in the two columns (reward relevant and irrelevant saccades) are for equal peak velocity movements. **(F)** Similar to part E, but for the group of mossy fibers that encoded motor commands with a delay. Error bars are SEM.

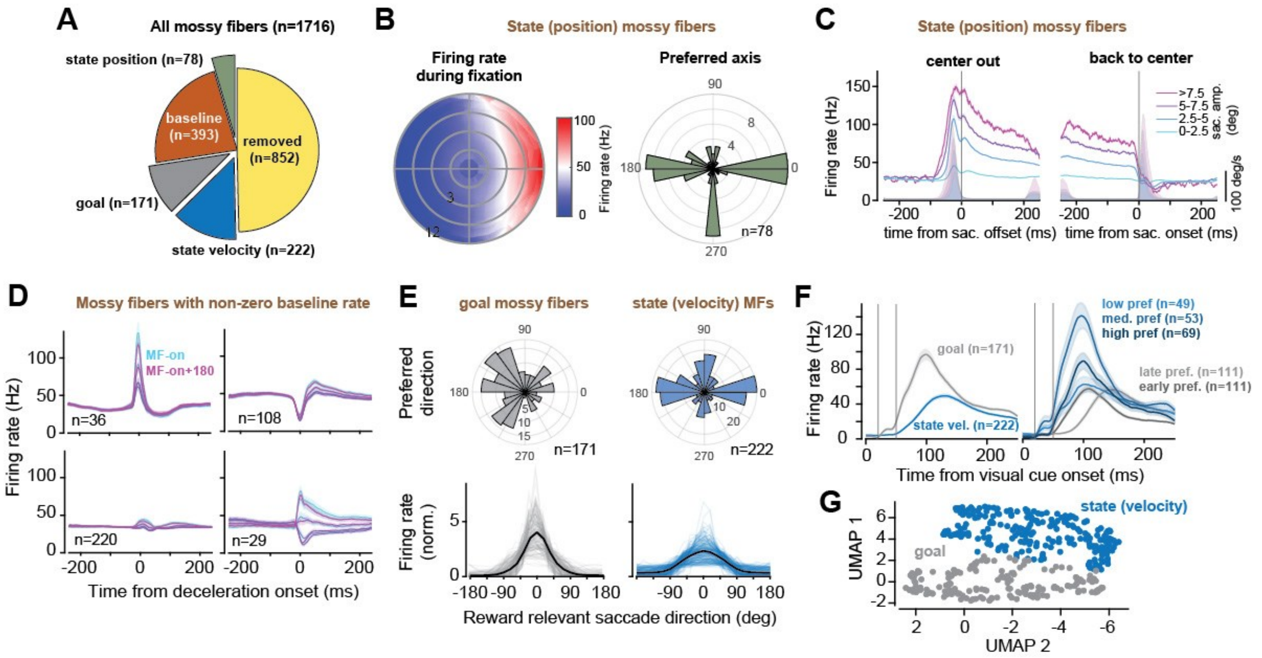


**Fig. S13. The properties of the mossy fibers.** (A) We recorded from n=1716 MFs. Roughly half were removed because the number of trials was less than 500 relevant saccades toward their preferred direction and less than 70 Hz of change in their response aligned to deceleration onset of the saccades. **(B)** State MFs that encoded position of the eyes. Left plot shows firing rates as a function of location of fixation aligned to the neuron's preferred axis. Right plot shows distribution of preferred axes. (C) Firing rates of state MFs that encoded position of the eyes, shown for saccades away from the center and then back to center. **(D)** MFs that had non-zero baseline activity, but the baseline rates did not vary significantly with position of the eyes. **(E)** Top: Distribution of preferred direction of the goal and state mossy fibers. The state mossy fibers encode the movements only along the horizontal and vertical axes, consistent with activity of burst generators in the brainstem. Bottom: firing rates in the goal and state MFs as a function of saccade direction with respect to preferred direction. State MFs have a broader width of direction encoding. **(F)** Response of goal and state MFs to visual cue. Some goal MFs exhibited a visual response but not the state MFs. **(G)** Goal and state MF clustering based on their response to visual target and motor responses. To make this map, we used a normalized distance measure between the responses to reward relevant and irrelevant saccades for similar saccade kinematics (direction, amplitude, peak velocity). In addition, we use the norm2 distance between the polar firing rate heat maps in the relevant and irrelevant saccades (for example, center heat maps in Fig. S11).

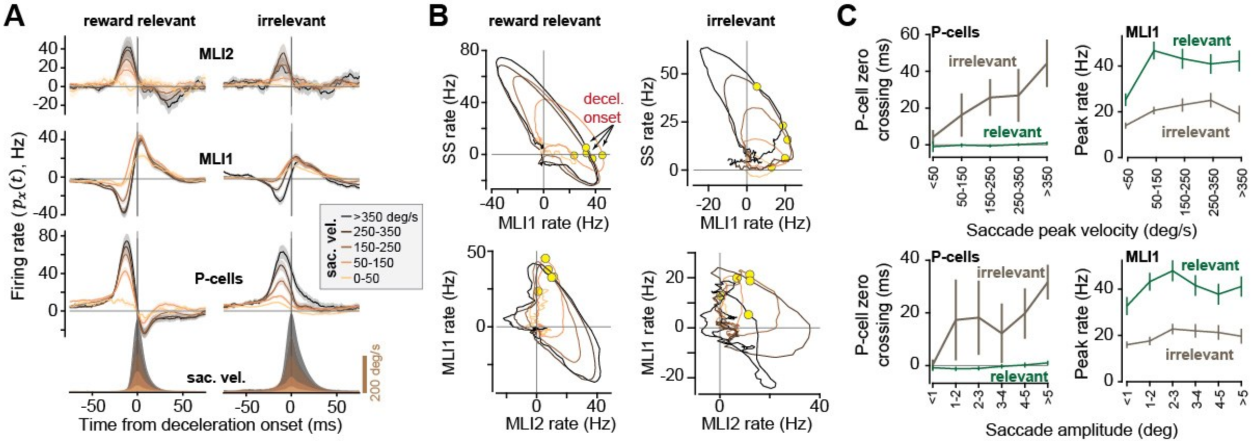


**Fig. S14**. **Population responses of the P-cells and pMLIs projected onto the direction of the movement for the two groups of saccades (reward relevant and irrelevant) binned by peak velocity of saccades. (A)** (left) pMLIs and P-cells population response binned by peak saccade velocity. (right) Same as left for the irrelevant saccades. **(B)** (top) Phase plots of P-cell responses versus pMLI1 responses for the two types of saccades. (bottom) Same as (top) for pMLI1s versus pMLI2s. In all cases, the relationship between MLI1 rates and SS rates has a negative slope, which indicates that an increase in the pMLI1 rate was associated with a decrease in the SS rates. This negative correlation is also present between pMLI2 and pMLI1. **(C)** P-cell zero-crossing and pMLI1 peak rate for different saccade (top) peak velocities (bottom) amplitudes.

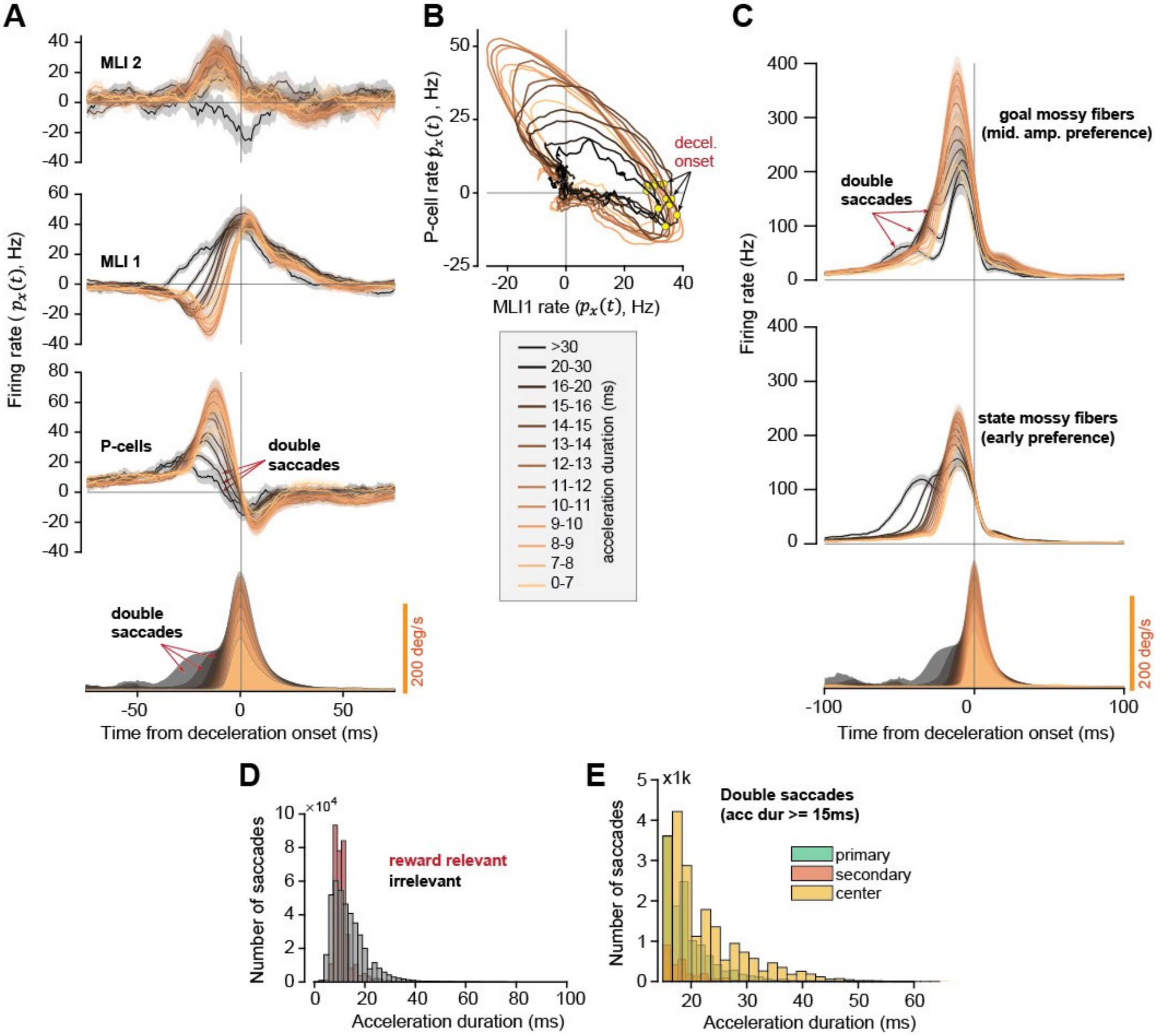


**Fig. S15**. **Population response of P-cells transitioned from burst to pause at deceleration onset, except for double saccades.** This may be because during double saccades there was a change in the goal information that the cerebellum received through the mossy fibers. **(A)** P-cells and pMLIs population response for reward relevant saccades projected onto the direction of the saccade, binned by acceleration duration of the saccade. Saccades with very long acceleration durations (>15ms) show a transient pause in their velocities, indicative of double saccades. **(B)** P-cells and pMLI1s remain on a similar manifold for all acceleration durations, including double saccades, implying that there is always a negative correlation between rate changes in pMLI1s and P-cells. **(C)** Response of goal and state mossy fibers as a function of saccade acceleration duration. Note the double peaked response in the mossy fibers for the double saccades. **(D)** Number of double saccades in the reward relevant and irrelevant saccades. **(E)** Double saccades were detected by acceleration durations longer than 15 ms and were present in all saccades, including primary, secondary, and center saccades.

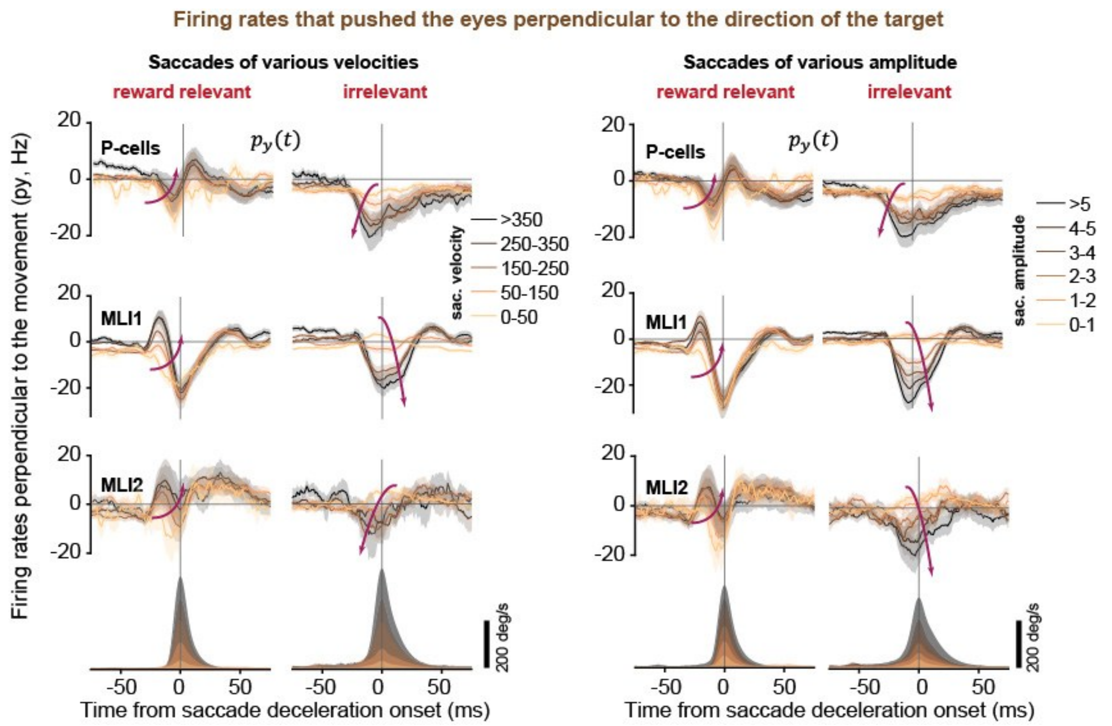


**Fig. S16**. **Cancellation of spikes that would move the eyes perpendicular to the direction of the target improved with saccade amplitude and velocity, but only if the movement was reward relevant.** The plots show the firing rates in the population of P-cells, MLI1s, and MLI2s, perpendicular to the direction of the movement. The left two columns display rates as a function of saccade velocity, comparing reward relevant and irrelevant saccades. Despite the fact that with increased velocity the neurons exhibit increased spike rates, a greater fraction of those rates is cancelled. However, this pattern does not hold for reward-irrelevant saccades. The right two columns display rates as a function of saccade amplitude. Error bars are SEM.

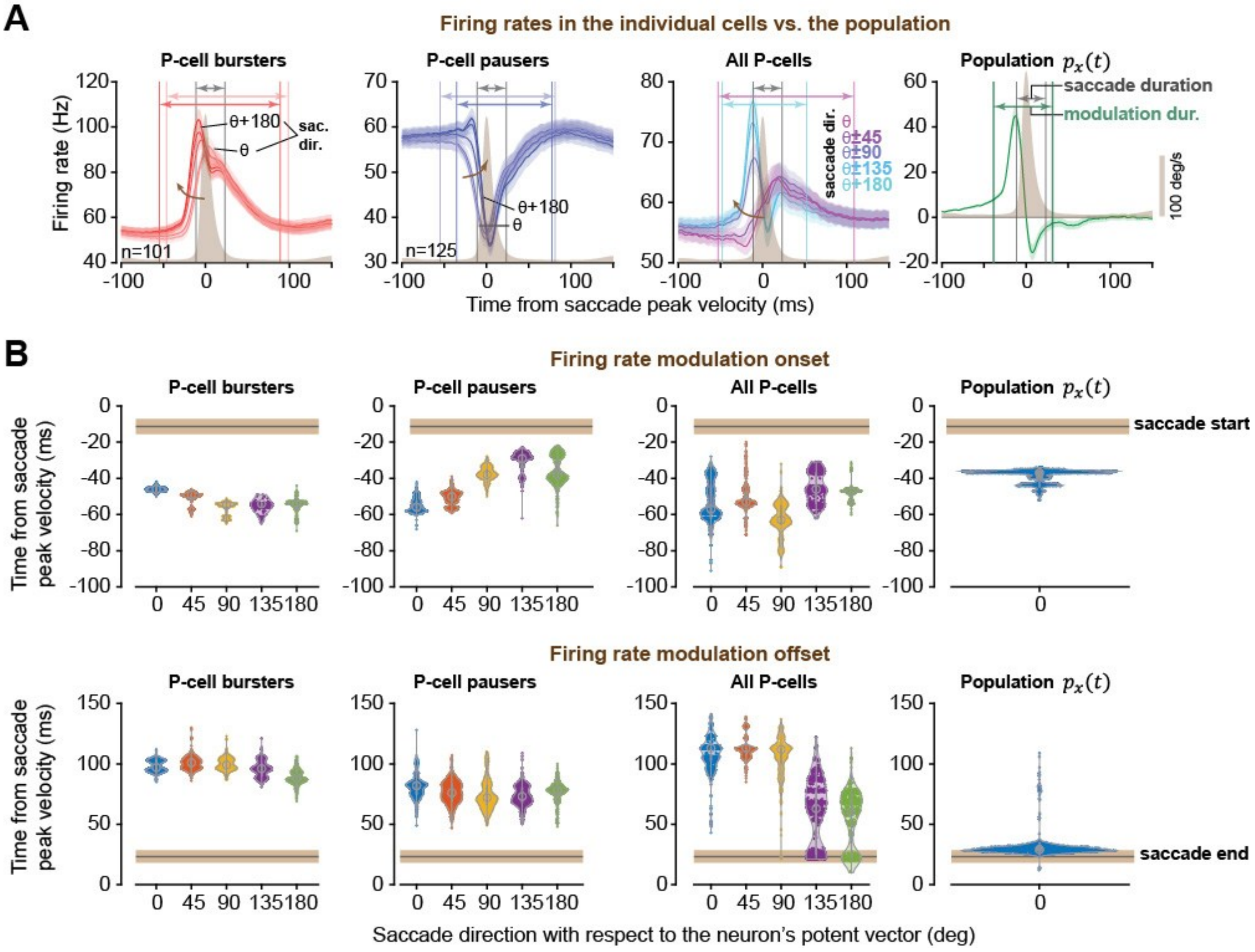


**Fig. S17**. **Spike cancellation among the P-cells allowed for production of a fast-changing output that ended as the movement ended. In contrast, individual P-cells exhibited firing rate modulations that changed slowly, returning to baseline much later than movement completion. (A)** Firing rates of individual P-cells and the population. The rates for the pausers and bursters are shown as a function of saccade direction with respect to the direction of the neuron's potent vector $\theta$. In the population plot, the response along the potent vector is plotted. **(B)** Timing of firing rate modulation onset and offset among the P-cells and in the population. Top row shows the timing of modulation onset. Bottom row shows the timing of the modulation offset. The horizontal bar indicating saccade start and end is shown with SD error bars. All error bars are SEM.

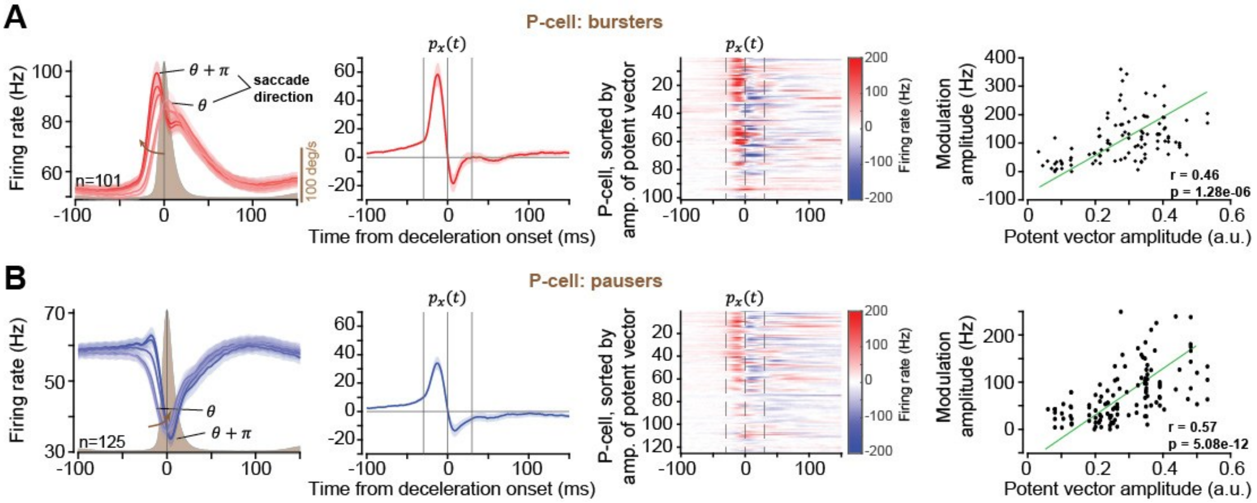


**Fig. S18**. **The magnitude of the burst-pause pattern in the population response of the P-cells was correlated to the size of their potent vector. (A)** P-cell bursters. From left to right: 1. The firing rates for saccades at various directions with respect to the potent vector. 2. The rate difference along the potent vector. 3. The rate difference along the potent vector, exhibited for each P-cell burster, sorted by the amplitude of its potent vector. 4. The relationship between the amplitude of the potent vector and the magnitude of the burst-pause. The magnitude was computed for each P-cell by subtracting the peak of activity during the acceleration phase (-30 to 0ms) minus trough during deceleration phase (0 to 30ms) (with respect to peak speed). **(B)** Same as in part A, but for P-cell pausers. All error bars are SEM.

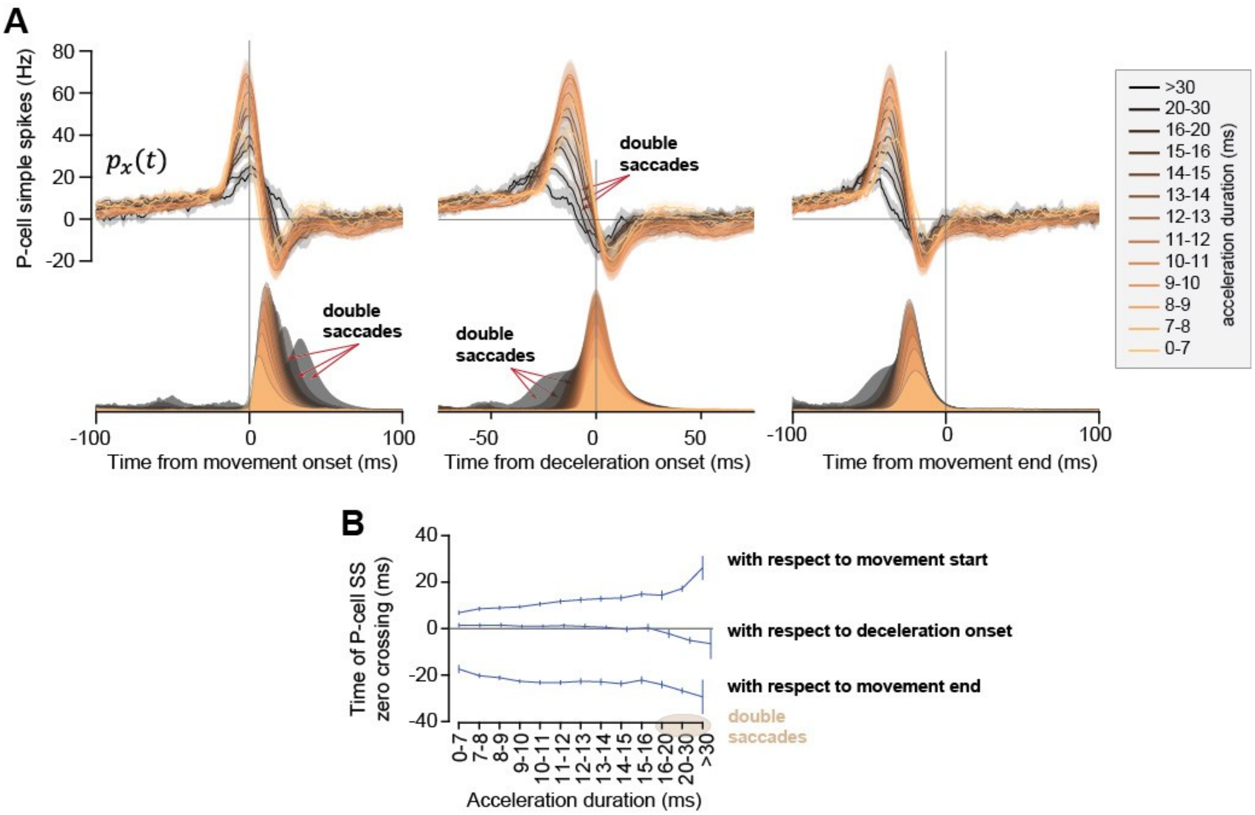


**Fig. S19**. **The P-cell simple spikes in the population response produced a zero crossing that remained time-locked to the onset of movement deceleration. (A)** P-cell population response aligned to movement onset, deceleration onset, and movement end. Double-saccades, i.e., saccades for which the subject appeared to change the goal mid-flight, are noted. **(B)** Timing of the zero crossing in the simple spike population response as a function of acceleration duration.

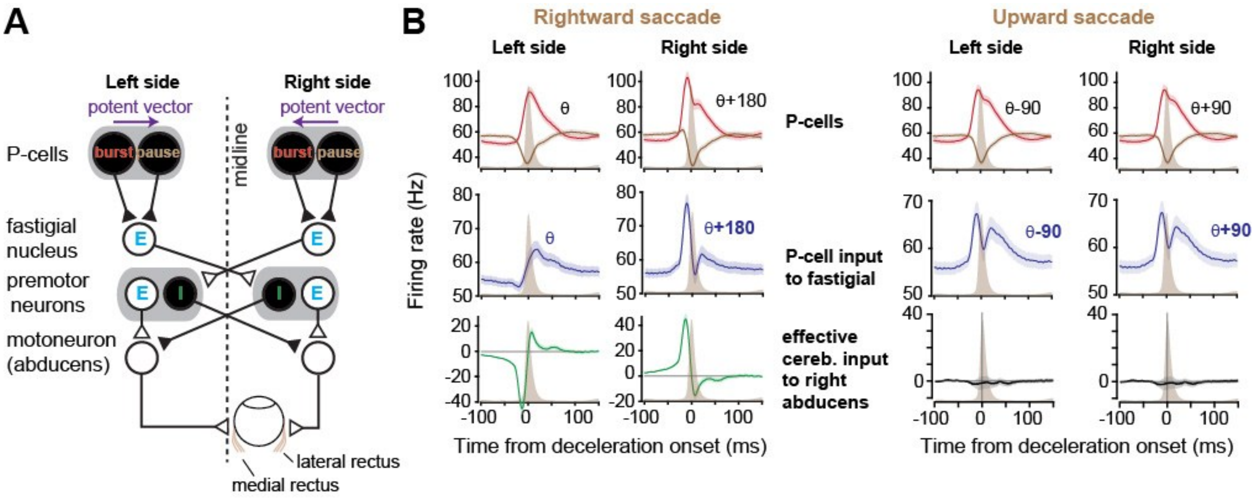


**Fig. S20. Anatomy of spike cancellation from the P-cells to the abducens nucleus: a model. (A)** Consider two groups of P-cells, one on the right side of the vermis, and one on the left. E: excitatory, I: inhibitory neuron. **(B)** A rightward horizontal saccade is against the potent vector for the P-cells on the right side of the vermis. That is, suppression of these neurons produces a displacement along their potent vector. As a result, optogenetic stimulation of these P-cells produces acceleration of rightward saccades (*53*). The same movement is along the potent vector for the P-cells on the left side. On the right side, the sum of P-cell activities (bursters and pausers) produces an early burst of inhibition onto the nucleus neurons. On the left side, the sum is a late burst of inhibition. Thus, fastigial activity is late for ipsilateral saccades, but early for contralateral saccades (*54*–*57*). The output from the fastigial reaches the contralateral burst generators (*58*), which are composed of inhibitory and excitatory neurons. The inhibitory burst generators (IBNs) project onto the contralateral abducens neurons, while the excitatory burst generators (EBNs) converge onto the ipsilateral abducens (*59*). As a result, for the rightward saccade, the effect of cerebellar output is to assist the eyes by exciting the right lateral rectus early, then stop the eyes by activating the medial rectus late into the movement (*60*–*62*). **(C)** During an upward (vertical saccade), the movement is 90 deg to the potent vector of the P-cells considered here. Thus, there needs to be complete cancellation of spikes that reach the motoneurons. This cancellation does not occur in the fastigial nucleus, as the neurons are active in all directions (*54*). It also does not occur in the IBNs or the EBNs, as both groups are active for vertical saccades (*63*–*65*). The P-cells produce precisely the same pattern of spikes on the right and the left side of the cerebellum for this movement. This produces converging activities onto the nucleus neurons that balance the excitation received by the burst generators on the left side with the excitation imposed on the burst generators on the right side. This balanced activation on the burst generators on the left and right (*66*) results in balanced excitation and inhibition at the motoneurons, resulting in cancellation (*67*). Thus, for this vertical movement, cancellation of the spikes that would have produced a horizontal displacement occurs not in the cerebellar nucleus, not in the premotor neurons (burst generators), but in the convergence of the inhibitory and excitatory inputs onto the motoneurons that pull the eyes in the horizontal direction.

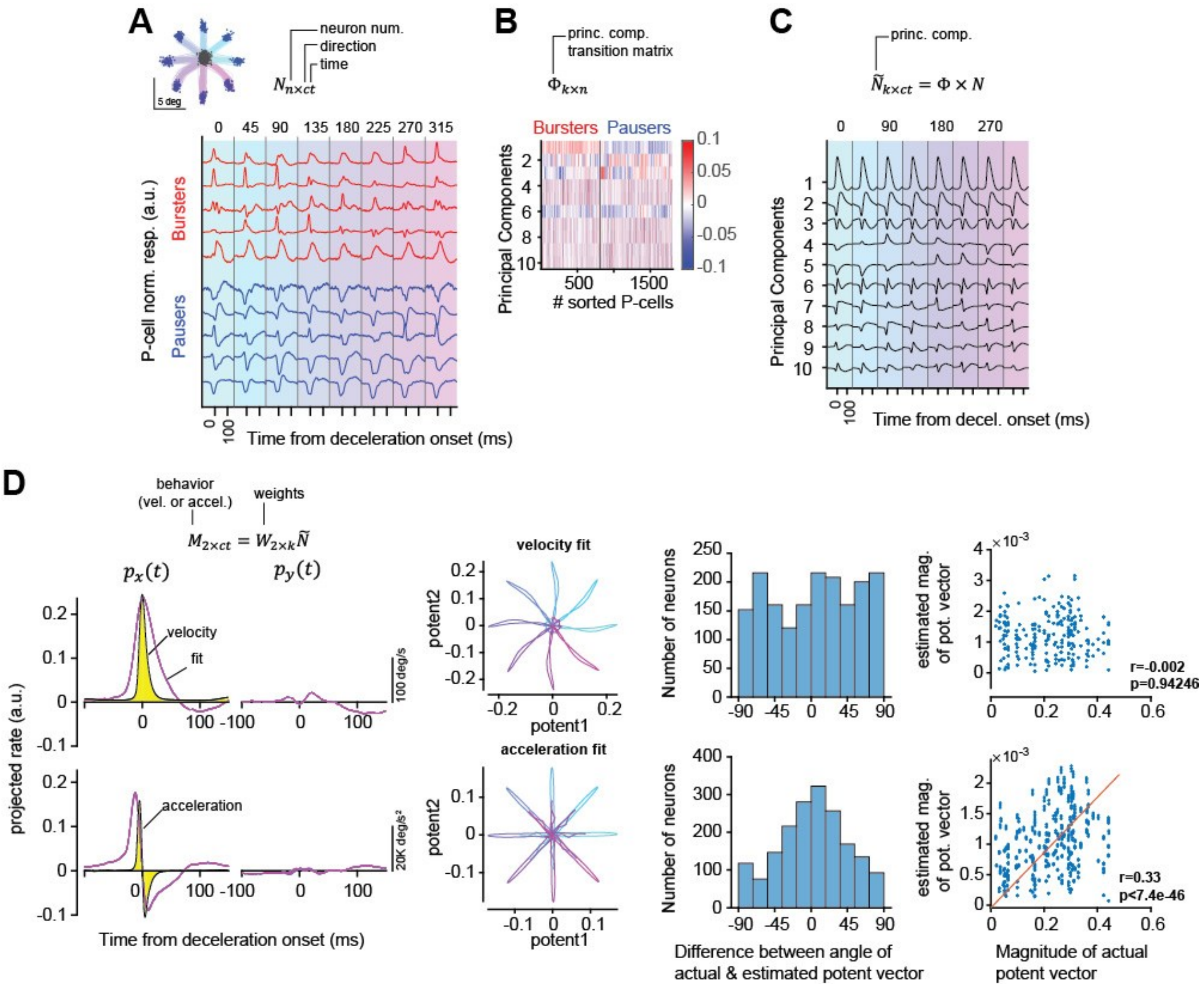


**Fig. S21. Comparing traditional approaches in finding potent vectors with the potent vector inferred from the climbing fiber inputs. (A)** The matrix $N$ represents the measured activities of P-cells (simple spikes), aligned to saccade deceleration onset, for each direction. This matrix was formed by assuming an unbiased sampling of the anatomical distribution of P-cells, i.e., a uniform distribution of climbing fiber indicated preferred directions. **(B)** The matrix $\Phi$, containing the first 10 principal components of $N$. **(C)** Matrix $\tilde{N}$. **(D)** We found the potent vector for each neuron by mapping $\tilde{N}$ onto behavior. For eye movements, a proxy for EMG is eye velocity because the burst generators that project onto the motoneurons encode eye velocity along either the horizontal or vertical axes. The top row shows the results of mapping the neural activity onto eye velocity. The second row shows the results if we assume that acceleration is the output that is being computed by the network. The first and second columns are the results of the fit along the target direction and perpendicular to target direction. The 4$^{th}$ and 5$^{th}$ columns are the relationship between the $W$ found using this approach, and the potent vector found from the climbing fiber inputs. Compare the results in the first column to the results from the climbing fiber derived potent vector in the right column of Fig. S19A.
